# SETH predicts nuances of residue disorder from protein embeddings

**DOI:** 10.1101/2022.06.23.497276

**Authors:** Dagmar Ilzhoefer, Michael Heinzinger, Burkhard Rost

**Affiliations:** Faculty of Informatics, Bioinformatics & Computational Biology, TUM (Technical University of Munich), Garching/Munich, Germany; Center of Doctoral Studies in Informatics and its Applications (CeDoSIA), TUM Graduate School, Garching, Germany; Institute for Advanced Study (TUM-IAS), TUM (Technical University of Munich), Garching/Munich, Germany; TUM School of Life Sciences Weihenstephan (WZW), TUM (Technical University of Munich), Freising, Germany

**Keywords:** protein disorder, residue disorder, IDP, IDR, protein structure prediction, AlphaFold2, protein language model

## Abstract

Predictions for millions of protein three-dimensional structures are only a few clicks away since the release of *AlphaFold2* results for UniProt. However, many proteins have so-called intrinsically disordered regions (IDRs) that do not adopt unique structures in isolation. These IDRs are associated with several diseases, including Alzheimer’s Disease. We showed that three recent disorder measures of *AlphaFold2* predictions (pLDDT, “experimentally resolved” prediction and “relative solvent accessibility”) correlated to some extent with IDRs. However, expert methods predict IDRs more reliably by combining complex machine learning models with expert-crafted input features and evolutionary information from multiple sequence alignments (MSAs). MSAs are not always available, especially for IDRs, and are computationally expensive to generate, limiting the scalability of the associated tools. Here, we present the novel method SETH that predicts residue disorder from embeddings generated by the protein Language Model ProtT5, which explicitly only uses single sequences as input. Thereby, our method, relying on a relatively shallow convolutional neural network, outperformed much more complex solutions while being much faster, allowing to create predictions for the human proteome in about one hour on a consumer-grade PC with one NVIDIA GeForce RTX 3060. Trained on a continuous disorder scale (CheZOD scores), our method captured subtle variations in disorder, thereby providing important information beyond the binary classification of most methods. High performance paired with speed revealed that SETH’s nuanced disorder predictions for entire proteomes capture aspects of the evolution of organisms. Additionally, SETH could also be used to filter out regions or proteins with probable low-quality *AlphaFold2* 3D structures to prioritize running the compute-intensive predictions for large data sets. SETH is freely publicly available at: https://github.com/Rostlab/SETH.

## 1 Introduction

### IDRs crucial for life

Protein sequence determines protein three-dimensional (3D) structure, which, in turn, determines protein function. While this dogma usually refers to proteins folding into well-defined 3D structures, other proteins do not adopt unique 3D structures in isolation. Instead, these so-called <u>intrinsically disordered proteins (IDPs;</u> (Dunker et al., 2013)) with <u>intrinsically disordered regions (IDRs)</u> sample their accessible conformational space, thereby expanding their functional spectrum (Wright and Dyson, 1999;Radivojac et al., 2004;Tompa et al., 2005;Tompa et al., 2006;Tompa et al., 2008;Uversky et al., 2009;Schlessinger et al., 2011) and possibly providing mechanisms to cope with evolutionary challenges (Tantos et al., 2009;Vicedo et al., 2015a;Vicedo et al., 2015b). The difference between long IDRs and long loops (neither helix nor strand) can be reliably predicted from sequences (Schlessinger et al., 2007b). For very short regions, IDRs and loops are technically not distinguishable in a predictive sense. Therefore, IDRs have to be longer than some minimal length Lmin for identification. While the precise value for Lmin remains obscure, Lmin=10 is clearly too short and Lmin=30 is clearly sufficient, as may be many values in between (Schlessinger et al., 2011). Using the more conservative Lmin=30, about 20-50% of all proteins in an organism are predicted to contain IDRs, with higher abundance in eukaryotes, especially in mammals (Romero et al., 1998;Liu et al., 2002;Schlessinger et al., 2011). Additionally, every fourth protein has been predicted as completely disordered (Dunker et al., 2008). This ubiquitous nature of disorder highlights its importance for the correct functioning of cells and makes the identification of IDRs crucial for understanding protein function. Alzheimer’s disease and Huntington’s disease, which are related to malfunctioning of disordered proteins/IDRs upon mutation, further underline this importance (Dyson and Wright, 2005;Dunker et al., 2008).

### CheZOD scores best characterize IDRs experimentally

The experimental study of protein disorder remains difficult. X-ray crystallography is challenged by the lack of rigidity and nuclear magnetic resonance (NMR) remains limited to proteins shorter than average (~450 residues; (Howard, 1998;Oldfield et al., 2013;Nwanochie and Uversky, 2019)). An additional complication is that upon binding to substrates, IDRs may appear ordered (Nielsen and Mulder, 2019). Arguably, today’s best experimental approach toward capturing IDRs are NMR-derived chemical shift Z-scores (<u>CheZOD scores</u>), despite the length-limitation (Nielsen and Mulder, 2019). In contrast to binary measures such as “missing X-Ray coordinates” (Romero et al., 1998), CheZOD scores provide a well-calibrated measure for the nuances of per-residue disorder. CheZOD scores are computed from the difference of chemical shift values obtained in NMR spectroscopy (Howard, 1998) and computed random coil chemical shift values (Nielsen and Mulder, 2020).

### Many prediction methods available

The limited scalability of labor-intensive and expensive wet-lab experiments has spawned many computational tools predicting IDRs, including (from old to new): PONDR (Romero et al., 1998;Peng et al., 2005), NORSp (Liu et al., 2002), DISOPRED2 (Ward et al., 2004), IUPred (Dosztanyi et al., 2005), FoldIndex (Prilusky et al., 2005), RONN (Yang et al., 2005), PrDOS (Ishida and Kinoshita, 2007), NORSnet (Schlessinger et al., 2007a), PreDisorder (Deng et al., 2009), MetaDisorder-MD (Schlessinger et al., 2009), ESpritz (Walsh et al., 2012), MetaDisorder (Kozlowski and Bujnicki, 2012), AUCpreD (Wang et al., 2016), SPOT-Disorder (Hanson et al., 2016), SPOT-Disorder-Single (Hanson et al., 2018), SPOT-Disorder2 (Hanson et al., 2019), rawMSA (Mirabello and Wallner, 2019), ODiNPred (Dass et al., 2020) and flDPnn (Hu et al., 2021). As for almost every phenotype since the introduction of the combination of machine learning and evolutionary information (<u>EI</u>), derived from multiple sequence alignments (MSAs; (Rost and Sander, 1993)), MSA-based predictions out-performed methods not using MSAs (Nielsen and Mulder, 2019;Dass et al., 2020). However, using MSAs slows down inference and performs worse for proteins in small families. This complicates the prediction of IDRs, which are inherently difficult to align due to, e.g., reduced sequence conservation in comparison to structured regions (Radivojac et al., 2002;Lange et al., 2016).

Besides these methods directly predicting disorder, *AlphaFold2* (Jumper et al., 2021), Nature’s method of the year 2021 (Marx, 2022), which provided a leap in the quality of protein structure predictions from MSAs and increases the width of structural coverage (Bordin et al., 2022), also provides measures indicative of IDRs. One of these, the pLDDT (predicted local distance difference test), estimates the performance of *AlphaFold2* depending on prediction strength, i.e., it measures prediction reliability as introduced for secondary structure prediction (Rost and Sander, 1993). However, instead of measuring it from the class output, *AlphaFold2* uses different objective functions and predicts its own reliability. The pLDDT distinguishes formidably well between trustworthy and less reliable predictions (Jumper et al., 2021). Additionally, low values for pLDDT have been suggested to predict IDRs rather accurately (Akdel et al., 2021;Wilson et al., 2021;Piovesan et al., 2022) or to predict non-existing proteins (Monzon et al., 2022). Furthermore, the “experimentally resolved” prediction of *AlphaFold2* should also contain information on disorder, since missing coordinates in experimentally recorded structures were an established definition of disorder (Dunker et al., 1998;Monastyrskyy et al., 2014). Lastly, the relative solvent accessible surface area of a residue (RSA (Connolly, 1983;Rost and Sander, 1994)) and its window average, calculated for *AlphaFold2* structure predictions, were also reported to be disorder predictors (Akdel et al., 2021;Piovesan et al., 2022;Redl et al., 2022).

Here, we bypassed the problem of generating MSAs for IDRs, by using embeddings from pre-trained protein language models (pLMs). Inspired by recent leaps in Natural Language Processing (NLP), pLMs learn to predict masked amino acids (tokens) given their surrounding protein sequence (Asgari and Mofrad, 2015;Alley et al., 2019;Bepler and Berger, 2019;Heinzinger et al., 2019;Bepler and Berger, 2021;Elnaggar et al., 2021;Ofer et al., 2021;Rives et al., 2021;Wu et al., 2021). Toward this end, amino acids correspond to words/tokens in NLP, while sentences correspond to full-length proteins in most current pLMs. As no information other than the amino acid sequence is required at any stage (*self-supervised learning*), pLMs efficiently leverage large but unlabeled databases with billions of protein sequences, such as BFD with more than 2 billion sequences (Steinegger et al., 2019). The information learned by the pLM during so-called (pre-) training can be retrieved and transferred afterwards (*transfer learning*), by encoding a protein sequence in vector representations (<u>embeddings</u>). In their simplest form, embeddings mirror the last “hidden” states/values of pLMs. In analogy to NLPs implicitly learning grammar, embeddings from pLMs capture some aspects of the language of life as written in protein sequences (Alley et al., 2019;Heinzinger et al., 2019;Ofer et al., 2021;Rives et al., 2021), which suffices as exclusive input to many methods predicting aspects of protein structure and function (Asgari and Mofrad, 2015;Alley et al., 2019;Heinzinger et al., 2019;Elnaggar et al., 2021;Heinzinger et al., 2021;Littmann et al., 2021a;Littmann et al., 2021b;Littmann et al., 2021c;Marquet et al., 2021;Rives et al., 2021).

First, we compared to which extent embeddings from five pLMs (ESM-1b (Rives et al., 2021), ProtBERT (Elnaggar et al., 2021), SeqVec (Heinzinger et al., 2019), ProtT5 (Elnaggar et al., 2021) and ProSE (Bepler and Berger, 2021)) could predict the degree of disorder of a residue as defined by CheZOD scores. Toward that end, we fit a minimal machine learning model (linear regression) on each of the five pLM embeddings. No pLM was fine-tuned in any way. The best performing embeddings served as input to partly a little more complex models, namely a logistic regression (LogReg), another linear regression (LinReg; trained on the full training set, as opposed to the linear regression used to compare pLMs which was only trained on 90% of the training set), a two-layer neural network (ANN), and a two-layer convolutional neural network (CNN; dubbed <u>SETH</u> (**S**elf-supervised **E**mbeddings predic**T** chemical s**H**ift Z-scores)). By training regression and classification models, we also investigated the benefit of training on nuanced CheZOD scores compared to binary disorder classification. The combination of using a rather simplistic model and embeddings from single protein sequences enabled the final method SETH to predict disorder for the entire Swiss-Prot with over 566,000 proteins (The UniProt et al., 2021) in approximately 7 hours on a machine with one RTX A6000 GPU with 48GB vRAM.

Since recent work showed that *AlphaFold2’*s (smoothed) pLDDT and (smoothed) RSA can be used to predict disorder (Akdel et al., 2021;Wilson et al., 2021;Piovesan et al., 2022;Redl et al., 2022), we tested *AlphaFold2* on CheZOD scores (following the advice of John Jumper, we also analyzed “experimentally resolved” predictions). Furthermore, we investigated the agreement between the disorder predictions of our best method and the pLDDT for 17 organisms, to establish SETH as a speed-up pre-filter for *AlphaFold2*. Lastly, we visually analyzed whether the predicted disorder spectrum carried any information about the evolution of 37 organisms.

## 2 Methods

### 2.1 Data sets

#### CheZOD scores

To streamline comparability to existing methods, we used two datasets available from ODiNPred (Dass et al., 2020) for training (file name CheZOD1325 in GitHub; 1325 proteins) and testing (file name CheZOD in GitHub; 117 proteins). Each dataset contains protein sequences and CheZOD scores for each residue. The CheZOD score measures the degree of disorder of the residue and is calculated from the difference between chemical shift values obtained by NMR spectroscopy (Howard, 1998) and computed random coil chemical shifts (Nielsen and Mulder, 2020). These differences vary considerably between ordered and disordered residues, thereby continuously measuring the nuances of order/ disorder for each residue (Nielsen and Mulder, 2020).

#### Redundancy reduction (*CheZOD1174* and *CheZOD117*)

To avoid overestimating performance through pairs of proteins with too similar sequences between training and testing sets, we constructed non-redundant subsets. Firstly, we built profiles (position specific scoring matrices; PSSMs) from multiple sequence alignments (MSAs) for proteins in the test set, obtained through three iterations with *MMSeqs2* ((Steinegger and Söding, 2017); using default parameters, except for “--num-iterations 3”, an established number of iterations, also applied in ColabFold (Mirdita et al., 2022) and enabling sensitive but still fast sequence searches (Steinegger and Söding, 2017)) against proteins in the training set. Next, any protein in the training set with >20% PIDE (percentage pairwise sequence identity) to any test set profile using bi-directional coverage (with default coverage threshold of 80%, focusing on joining proteins with similar domain composition (Hauser et al., 2016)) was removed using *MMSeqs2* high-sensitivity (--s 7.5) search. The value PIDE<20% was, for simplicity, concluded from an earlier analysis of the reach of homology-based inference for the structural similarity of protein pairs (Rost, 1999). The training set had been constructed such that all protein pairs had <50% PIDE (Dass et al., 2020), and we did not reduce the redundancy within the training set any further. Secondly, we removed all residues without valid CheZOD scores (indicated by CheZOD scores≥900; for all models apart from SETH, they were removed after embedding generation, while for SETH (CNN) they were removed before, to enable undisturbed passing of information from neighboring residues). The resulting training set (dubbed <u>*CheZOD1174*</u>) contained 1174 proteins with a total of 132,545 residues (at an average length of 113 residues, these proteins were about 3-4 times shorter than most existing proteins). The resulting dataset for testing (dubbed <u>*CheZOD117*</u>) contained 117 sequences with a total of 13,069 residues (average length 112). Consequently, we did not alter the test set published alongside ODiNPred, which has been used to evaluate 26 disorder prediction methods (Nielsen and Mulder, 2019), enabling a direct comparison of the results. However, we altered the training data published and used for ODiNPred, to reduce the overlap between training and testing.

#### Dataset distributions

After preparing the data, we analyzed the distributions of the CheZOD scores for both *CheZOD117* and *CheZOD1174* (Fig. S1). The CheZOD scores in these sets ranged from −5.6 to 16.2. Nielsen and Mulder had previously established a threshold of eight to differentiate between disorder (CheZOD score≤8) and order (CheZOD score>8) (Nielsen and Mulder, 2016). In both sets, the CheZOD score distributions were bimodal, but while there was an over-representation of ordered residues in the training set *CheZOD1174* (72% ordered), disordered residues were most prevalent in the test set *CheZOD117* (31% ordered). As artificial intelligence (AI) always optimizes for similar distributions in train and test, the train-test set discrepancy provided an additional safeguard against over-estimating performance.

### 2.2 Input embeddings

#### Five pLMs

Protein sequences from both sets (*CheZOD117*, *CheZOD1174*) were encoded as distributed vector representations (embeddings) using five pLMs: (1) SeqVec (Heinzinger et al., 2019), based on the NLP algorithm ELMo (Peters et al., 2018), is a stack of bi-directional long short-term memory cells (LSTM (Hochreiter and Schmidhuber, 1997)) trained on a 50% non-redundant version of UniProt (The UniProt et al., 2021) (UniRef50 (Suzek et al., 2015)). (2) ProtBERT (Elnaggar et al., 2021), based on the NLP algorithm BERT (Devlin et al., 2018), trained on BFD, the Big Fantastic Database, (Steinegger and Söding, 2018;Steinegger et al., 2019) with over 2.1 billion protein sequences. (3) ESM-1b (Rives et al., 2021), conceptually similar to (Prot)BERT (both use a stack of Transformer encoder modules (Vaswani et al., 2017)), but trained on UniRef50. (4) ProtT5-XL-U50 (Elnaggar et al., 2021) (dubbed <u>ProtT5</u> for simplicity), based on the NLP sequence-to-sequence model T5 (Transformer encoder-decoder architecture) (Raffel et al., 2020), trained on BFD and fine-tuned on Uniref50. (5) ProSE (Bepler and Berger, 2021), consisting of LSTMs trained on 76M unlabeled protein sequences in UniRef90 and additionally on predicting intra-residue contacts and structural similarity from 28k SCOPe proteins (Fox et al., 2014). While ProtBERT and ESM-1b were trained on reconstructing corrupted tokens/amino acids from non-corrupted (protein) sequence context (i.e., masked language modeling), ProtT5 was trained by teacher forcing, i.e., input and targets were fed to the model, with inputs being corrupted protein sequences and targets being identical to the inputs but shifted to the right (span generation with span size of 1 for ProtT5). In contrast, SeqVec was trained on predicting the next amino acid, given all previous amino acids in the protein sequence (referred to as auto-regressive pre-training). All pLMs, except for ProSE, were optimized only through self-supervised learning, i.e., exclusively using unlabeled sequences for pre-training. In contrast, ProSE was trained on three tasks simultaneously (multi-task learning), i.e., masked language modeling was used to train on 76M unlabeled sequences in UniRef90 and training to predict residue-residue contacts together with structural similarity was performed using 28k labeled protein sequences from SCOPe (Fox et al., 2014).

#### Embeddings: last hidden layer

Embeddings were extracted from the last hidden layer of the pLMs, with ProtT5 per-residue embeddings being derived from the last attention layer of the model’s encoder-side using half-precision. The *bio_embeddings* package was used to generate the embeddings (Dallago et al., 2021). The resulting output is a single vector for each input residue, yielding an LxN-dimensional matrix (L: protein length, N: embedding dimension; N=1024 for SeqVec/ProtBERT/ProtT5; N=1280 for ESM-1b; N=6165 for ProSE).

#### Choosing embeddings best suited for IDR prediction

To find the most informative pLM embeddings for predicting IDRs/CheZOD score residue disorder, we randomly chose 90% of the proteins in *CheZOD1174* and trained a linear regression model on each of the five pLM embeddings to predict continuous CheZOD scores. To simplify the comparison and “triangulation” of our results, we also compared these five embedding-based models to inputting the standard one-hot encodings (i.e., 20 instead of 1024/1280/6165 input units per residue). One-hot encodings represent each residue/sequence position by a 20-dimensional vector, for the 20 standard amino acids essentially contained in all proteins. Each position in the vector corresponds to one amino acid, i.e., the elements of the vector are binary: 1 for the position in the vector corresponding to the encoded amino acid, 0 otherwise. The special case “X” (unknown amino acid) was encoded in a 20-dimensional vector containing only 0s. The linear regressions were implemented with the *LinearRegression* module of scikit-learn (Pedregosa et al., 2011) with all parameters left at default values. We evaluated the models on the remaining 10% of *CheZOD1174* using the Spearman correlation coefficient (ρ; Eqn. 2) and the AUC (area under the receiver operating characteristic curve; Eqn. 3; see Methods section 2.5).

#### Unsupervised embedding analysis

Lastly, we analyzed the ProtT5 embeddings of *CheZOD117* in more detail by creating a t-distributed stochastic neighbor embedding (t-SNE; (van der Maaten and Hinton, 2008)) using scikit-learn (Pedregosa et al., 2011). PCA (principle component analysis (Wold et al., 1987)) initialized the t-SNE to enable higher reliability of the resulting structure (Kobak and Berens, 2019). Furthermore, following a rule of thumb previously established (Kobak and Berens, 2019), the perplexity was chosen at the high value of 130 (1% of the sample size) to emphasize the global data structure (Kobak and Berens, 2019) in order to identify putative clusters of order or disorder (defaults for all other parameters).

### 2.3 New disorder prediction methods

We optimized four models to predict disorder: (1) linear regression (dubbed <u>LinReg</u>), (2) multi-layer artificial neural network (dubbed <u>ANN</u>), (3) two-layer CNN (dubbed <u>SETH</u>) and (4) logistic regression (dubbed <u>LogReg</u>). The models used throughout this work were deliberately kept simple to gain speed and avoid over-fitting. Three of our models were trained on regression (LinReg, ANN and SETH), while LogReg was trained on discriminating disordered from ordered residues (binary classification: disorder: CheZOD score≤8, order: CheZOD score>8; (Nielsen and Mulder, 2016)).

SETH was implemented in PyTorch (Paszke et al., 2019) using *Conv2d* for the convolutional layers, MSELoss as loss function and Adam as optimizer (learning rate of 0.001), activating amsgrad (Reddi, 2018). Additionally, we padded to receive one output per residue and set all random seeds to 42 for reproducibility. Lastly, we randomly split *CheZOD1174* into training (90% of proteins: optimize weights) and validation (10%: for early-stopping after 10 epochs without improvement, hyper-parameter optimization: of the best performing models, we chose that with the most constraints (Fig. S3, red bar), resulting in a kernel size of (5,1), 28 output channels of the first convolutional layer, the activation function Tanh between the 2 convolutional layers and the weight decay parameter of 0.001 in the optimizer). Details for LinReg, ANN and LogReg are in Supplementary Material 1.1 and Fig. S2.

### 2.4 AlphaFold2

*AlphaFold2* (Jumper et al., 2021) predicts a reliability for each residue prediction, namely, the pLDDT. This score and its running average over a window of consecutive residues have been claimed to predict disorder (Akdel et al., 2021;Wilson et al., 2021;Piovesan et al., 2022;Redl et al., 2022). Another objective function predicted by AlphaFold2, namely, the “experimentally resolved” prediction (Jumper et al., 2021) is likely also informative, as missing coordinates in experimental structures have been used to define disorder (Dunker et al., 1998;Monastyrskyy et al., 2014). To analyze these *AlphaFold2* predictions against CheZOD scores, we applied *ColabFold* (Mirdita et al., 2022) to predict 3D structures for all proteins in *CheZOD117*. *ColabFold* speeds up *AlphaFold2* predictions 40-60x mostly by replacing jackhmmer (Johnson et al., 2010) and HHblits (Remmert et al., 2012) in the computationally expensive MSA generation by *MMSeqs2* (Steinegger and Söding, 2017) without losing much in performance. We generated MSAs by searching UniClust30 (Mirdita et al., 2017) and the environment database ColabFoldDB (Mirdita et al., 2022). We used neither templates nor Amber force-field relaxation (Hornak et al., 2006), as those do not significantly improve performance (Jumper et al., 2021;Mirdita et al., 2022) although increasing runtime manifold (especially the Amber relaxation). As *ColabFold* currently does not support outputting the “experimentally resolved” head, we added this feature by averaging over the sigmoid of the raw “experimentally resolved” logits output of *AlphaFold2* for each atom in a residue. After having generated the pLDDT values and the “experimentally resolved” predictions, we additionally calculated the smoothed pLDDT for each residue, using a sliding window of 21 consecutive residues following previous findings (Akdel et al., 2021). While sliding the window over the sequence, the center residue of the window was always assigned the mean of all values within the window (instead of padding, windows closer than 10 residues to the N- and C-terminus were shrunk asymmetrically, e.g., for the N-terminal position i (i=1,..,10, starting at i=1 at the N-terminus): averaging over (i-1) positions left of i).

It has also been reported that the window-averaged RSA calculated from AlphaFold2’s 3D predictions correlates with IDRs (Akdel et al., 2021;Piovesan et al., 2022;Redl et al., 2022). Consequently, we also added this measure to our evaluation. With the pipeline provided alongside one of the analyses reporting the RSA as a disorder predictor (Piovesan et al., 2022), we calculated the RSA for the residues of the *CheZOD117* set, leaving all parameters at default. Then we smoothed the RSA by averaging over 25 consecutive residues as suggested elsewhere (Piovesan et al., 2022). While the use of the “experimentally resolved” predictions is new here (thanks to John Jumper for the recommendation), all other ways of processing *AlphaFold2* predictions to predict disorder were taken from other work.

### 2.5 Evaluation

We followed a previous analysis in evaluating our performance (same evaluation measures and test set) for easy comparability (Nielsen and Mulder, 2019). This allowed a direct comparison to (alphabetical list): AUCPred with and without evolution (Wang et al., 2016), DisEMBL (Linding et al., 2003), DISOPRED2 (Ward et al., 2004), DISOPRED3 (Jones and Cozzetto, 2015), DISpro (Cheng et al., 2005), DynaMine (Cilia et al., 2014), DISPROT/VSL2b (Vucetic et al., 2005), ESpritz (Walsh et al., 2012), GlobPlot (Linding et al., 2003), IUPred (Dosztanyi et al., 2005), MetaDisorder (Kozlowski and Bujnicki, 2012), MFDp2 (Mizianty et al., 2013), PrDOS (Ishida and Kinoshita, 2007), RONN (Yang et al., 2005), s2D (Sormanni et al., 2015), SPOT-Disorder (Hanson et al., 2016). We added results for flDPnn (Hu et al., 2021) and ODiNPred (Dass et al., 2020) using the publicly available web-servers. Unfortunately, we failed to accomplish the same for SPOT-Disorder2 ((Hanson et al., 2019); no server response).

We estimated the Spearman correlation, ρ, and its 95% confidence interval (CI) over n = 1000 bootstrap sets in all cases (Efron and Tibshirani, 1991). For each bootstrap set, a random sample of the size of the test set (=m) was drawn with replacement from the test set. For each of these sampled sets, the ρ was calculated. If u_i_ is the rank of the i^th^ value in the ground truth CheZOD scores and v_i_ the rank of the i^th^ value in the predicted CheZOD scores (or the rank of the respective predictive values for LogReg and *AlphaFold2*) of the method, the ρ was calculated with Eqn. 2. The final ρ was derived from averaging over those 1000 values and the 95% CI was estimated by computing the standard deviation of the ρ over the sampled sets and multiplying it by 1.96. The standard deviation was calculated with Eqn. 1, where x_i_ is the ρ of an individual bootstrap set and ⟨x⟩ is the average ρ over all bootstrap sets.

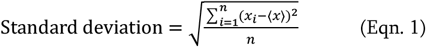

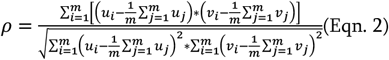

Furthermore, the AUC and its 95% CI were estimated for each model evaluated here, again, by applying the same bootstrapping procedure. As the AUC requires binarized ground truth class labels, continuous CheZOD scores were binarized using the threshold of eight (disorder CheZOD score≤8 and order CheZOD score>8; (Nielsen and Mulder, 2016)) for the calculation of the AUC (Eqn. 3; scikit-learn implementation). In Eqn. 3, I[.] is the indicator function, m^+/−^ are the number of ordered /disordered samples in the test set (classifying the samples according to the ground truth class label) and y_i_^+/−^ is the ith predicted value in the ordered /disordered samples.

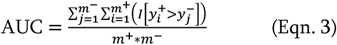

Lastly, we plotted the receiver operating characteristic curve for our models (SETH, LinReg/LinReg1D, ANN and LogReg), as well as for *AlphaFold2*’s pLDDT (Fig. S5).

### 2.6 Additional tests

#### Runtime

We analyzed the runtime for the best method introduced here (SETH), by clocking the predictions for the human proteome (20,352 proteins) and the Swiss-Prot database (566,969 proteins; (The UniProt et al., 2021)). This evaluation was performed on a machine with 2 AMD EPYC™ ROME 7352 CPUs at 2.30GHz each with 24/48 cores, a 256GB RAM (16 x 16GB) DDR4-3200MHz ECC, one RTX A6000 GPU with 48GB RAM, a 278GB SSD scratch disk and a 7.3TB HDD. However, the final model constituting SETH can also easily be deployed on any machine holding a GPU with ≥8GB RAM at some cost in speed, allowing to run SETH, e.g., in *Google Colab*. To reflect this, we also benchmarked the speed for running the entire human proteome on a smaller GPU (single NVIDIA GeForce RTX 3060 with 12GB vRAM). Lastly, we benchmarked the speed on our test set *CheZOD117* on an AMD Ryzen 5 5500U CPU, to reflect that SETH can even efficiently be run without a GPU for small sets. All values for runtime included all steps required: (1) load ProtT5, (2) load SETH model checkpoint, (3) read sequences from FASTA files, (4) create embeddings, (5) create predictions and (6) write results into a file.

#### Comparison: CheZOD score predictions and pLDDT in 17 organisms

From the AlphaFold database with 3D predictions (Jumper et al., 2021), we downloaded the available files ending in “F1-model-v2.pdb” for all proteins listed in UniProt (The UniProt et al., 2021) for 19 organisms (Table S2). A few files (0.3% of proteins) appeared with seemingly corrupted format (no separator between some values) and were removed. For all others, we extracted the pLDDT values.

For three organisms (*Leishmania infantum*, *Schistosoma mansoni* and *Plasmodium falciparum*) we predicted disorder with SETH using the sequences provided in UniProt (The UniProt et al., 2021); for the remaining 16 organisms (Table S2) we used the sequences present in Swiss-Prot, due to already having generated this data (see 2.6 Runtime). Due to GPU resources, we did not predict disorder for proteins with >9,000 residues (0% of *Leishmania infantum* + *Schistosoma mansoni*, 0.7% – 40 proteins – of *Plasmodium falciparum*, 0.004% – 25 proteins – of Swiss-Prot). None of the proteins for the CheZOD sets (CheZOD117, CheZOD1174) were that long (for obvious reasons related to the length-limitation of NMR).

To compare disorder predictions and pLDDT, only the subset of the data where both *AlphaFold2* and disorder predictions were available were used. The resulting set contained 17 of the above downloaded 19 organisms (Table S2; 2 organisms: no overlap in the predictions available for *AlphaFold2* and SETH) with 105,881 proteins containing a total of 47M residues. We referred to this data set as the *17-ORGANISM-set*.

#### Spectrum of predicted CheZOD score distributions for entire organisms

The spectra of predicted subcellular location reveal aspects pertaining to the evolution of species (Marot-Lassauzaie et al., 2021b). Consequently, we tried the same concept on predicted CheZOD scores for Swiss-Prot. For technical reasons (GPU memory), we excluded proteins longer than 9,000 residues from our analysis. In the entire Swiss-Prot, 0.004% of the proteins reached this length and were excluded. For the other 99.996%, we first converted all predicted CheZOD score distributions (consisting of all disorder predictions of all residues within one organism) of all Swiss-Prot organisms into vectors by counting CheZOD scores in eight bins (−15, −11.125, −7.25, −3.375, 0.5, 4.375, 8.25, 12.125, 16). After normalization (dividing raw counts by all residues in the organism), we PCA-plotted 37 organisms of Swiss-Prot with at least 1500 proteins ((Wold et al., 1987); to keep clarity in the plot, some organisms with at least 1500 proteins were neglected), using the standard implementation of R (prcomp; (Team, 2021)).

## 3 Results

### Success of minimalist: single sequence, simple model

While state-of-the-art (SOTA) methods usually rely on MSA input to predict IDRs, the methods introduced here use pLMs to encode single protein sequences as embeddings that served as the sole input feature for any prediction. To find the most informative pLM for IDRs, we predicted CheZOD scores through the minimalistic approach of linear regressions on top of embeddings from five pLMs (ProtT5 (Elnaggar et al., 2021), ProSE (Bepler and Berger, 2021), ESM-1b (Rives et al., 2021), ProtBERT (Elnaggar et al., 2021), SeqVec (Heinzinger et al., 2019)). Following the recent assessment of 26 methods (Nielsen and Mulder, 2019), we calculated the Spearman correlation coefficient ρ between true and predicted CheZOD scores and the AUC for 10% of *CheZOD1174*, not used in training, to evaluate the models.

Embeddings from all pLMs outperformed the random baseline and the one-hot encodings, both for the correlation (Fig. 1B; ρ) and the binary projection of CheZOD scores (Fig. 1A; AUC). The simplest pLM-type included here, namely *SeqVec*, performed consistently and statistically significantly worse than all other pLMs (Fig. 1). The other four embeddings (ProSE, ESM-1b, ProtT5, ProtBERT) did not differ to a statistically significant extent, given the small data set. However, since the linear regression trained on ProtT5 reached the numerical top both in ρ and AUC, we used only embeddings from ProtT5 for further analyses.

**Fig. 1:**
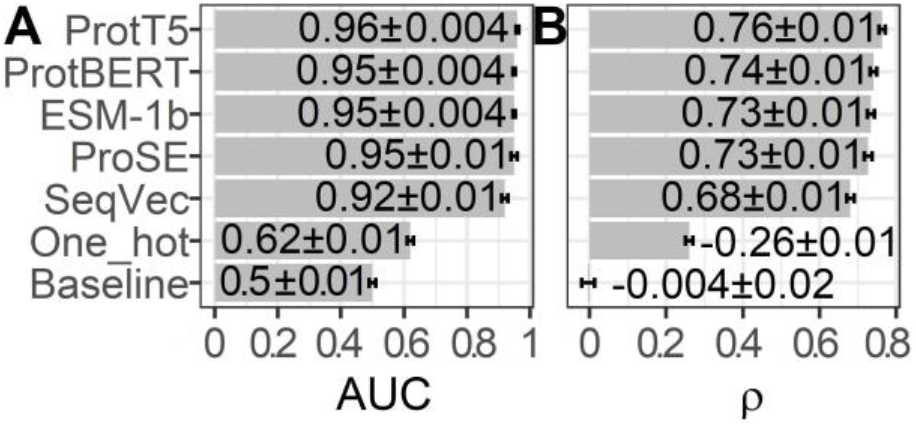
Linear regression on five pLMs sufficed. Performance estimates for training on 90% of CheZOD1174 (Dass et al., 2020) and testing on the remaining 10% using linear regressions fed by 20-dimensional one-hot encodings or raw embeddings (without further optimization) from five protein language models (pLMs): ProtT5 (Elnaggar et al., 2021), ProtBERT (Elnaggar et al., 2021), ESM-1b (Rives et al., 2021), ProSE (Bepler and Berger, 2021), SeqVec (Heinzinger et al., 2019). The seventh row displays the performance of the baseline/random model computed on 1024-dimensional embeddings sampled randomly from a standard normal distribution. Panel (A) required to first project predictions onto a binary state of disorder (CheZOD score≤8)/order (CheZOD score>8) and measures the area under the receiver operating characteristic curve (AUC; Eqn. 3), while Panel (B) depicts the Spearman correlation coefficient (ρ; Eqn. 2), calculated using the observed and predicted CheZOD scores. The errors mark the 95% confidence intervals approximated by multiplying 1.96 with the bootstrap standard deviation (Methods)

### ProtT5 captured disorder without any optimization

Next, we analyzed which information about disorder ProtT5 had already learned during self-supervised pre-training, i.e., before seeing any disorder-related labels. Towards this end, t-SNE projected the 1024-dimensional embeddings onto two dimensions (Fig. 2). This suggested some level of separation between ordered (red) and disordered (blue) residues (Fig. 2A: red colors oriented toward the center in each cluster), indicating that even raw ProtT5 embeddings already captured some aspects of disorder without seeing any such annotations (ProtT5 only learned to predict masked amino acid tokens (Elnaggar et al., 2021)). However, the major signal seemingly did not cluster the disorder/order phenotype. Instead, the primary 20 clusters corresponded to the 20 amino acids (Fig. 2B).

**Fig. 2:**
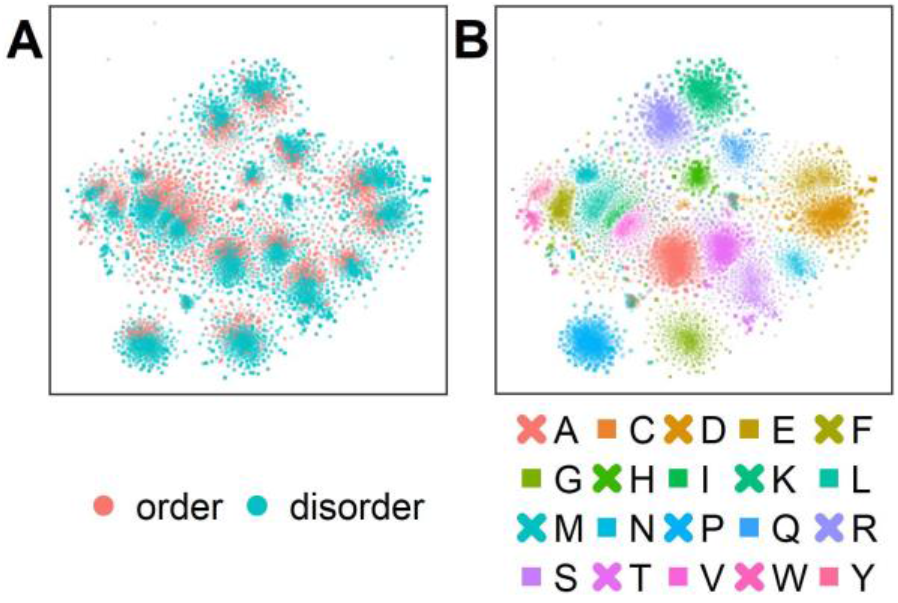
Raw embeddings carry some information about CheZOD score disorder. The t-SNE dimensionality reduction (van der Maaten and Hinton, 2008) was performed on the 1024-dimensional ProtT5 (Elnaggar et al., 2021) residue-level embeddings extracted from the last attention layer of ProtT5 for all sequences in test set CheZOD117 (13,069 residues; (Dass et al., 2020)). Panel (A) shows the embeddings colored by order (CheZOD score>8; red) and disorder (CheZOD score≤8, blue; (Nielsen and Mulder, 2016)). Panel (B) shows the same t-SNE projection but with coloring by the 20 standard amino acid types (here shown in one-letter code; A=Alanine, C=Cysteine, D=Aspartic acid, E=Glutamic acid, F=Phenylalanine, G=Glycine, H=Histidine, I=Isoleucine, K=Lysine, L=Leucine, M=Methionine, N=Asparagine, P=Proline, Q=Glutamine, R=Arginine, S=Serine, T=Threonine, V=Valine, W=Tryptophan, Y=Tyrosine).

### SETH (CNN) outperformed other supervised models

Next, we trained four AI models, inputting ProtT5 embeddings: three predicted continuous CheZOD scores (LinReg, ANN, SETH), one predicted binary disorder (LogReg). We could add the performance of our methods to a recent method comparison (Nielsen and Mulder, 2019) since we used the same performance metrics and test set (*CheZOD117;* Fig. 3). We also added the ODiNPred web application (Dass et al., 2020), the flDPnn webserver (Hu et al., 2021) and the performance of the new method ADOPT ESM-1b (Redl et al., 2022), which also uses pLM embeddings. When considering the mean ρ (Fig. 3A), our methods SETH and ANN numerically outperformed all others, both those not using MSAs (below dashed line in Fig. 3), and those using MSAs (above dashed line in Fig. 3). When requiring a statistically significant difference at the 95% CI (±1.96 standard errors) for the ρ, our methods (SETH, ANN, LinReg and LogReg) significantly outperformed all others, except for ODiNPred and ADOPT ESM-1b. When evaluating the performance based on the mean AUC, SETH and the simplistic LinReg outperformed all other evaluated methods. Due to the already high AUC levels of many methods, the absolute improvement of our models (SETH, ANN, LinReg and LogReg) to SOTA methods in terms of AUC was often not statistically significant.

**Fig. 3:**
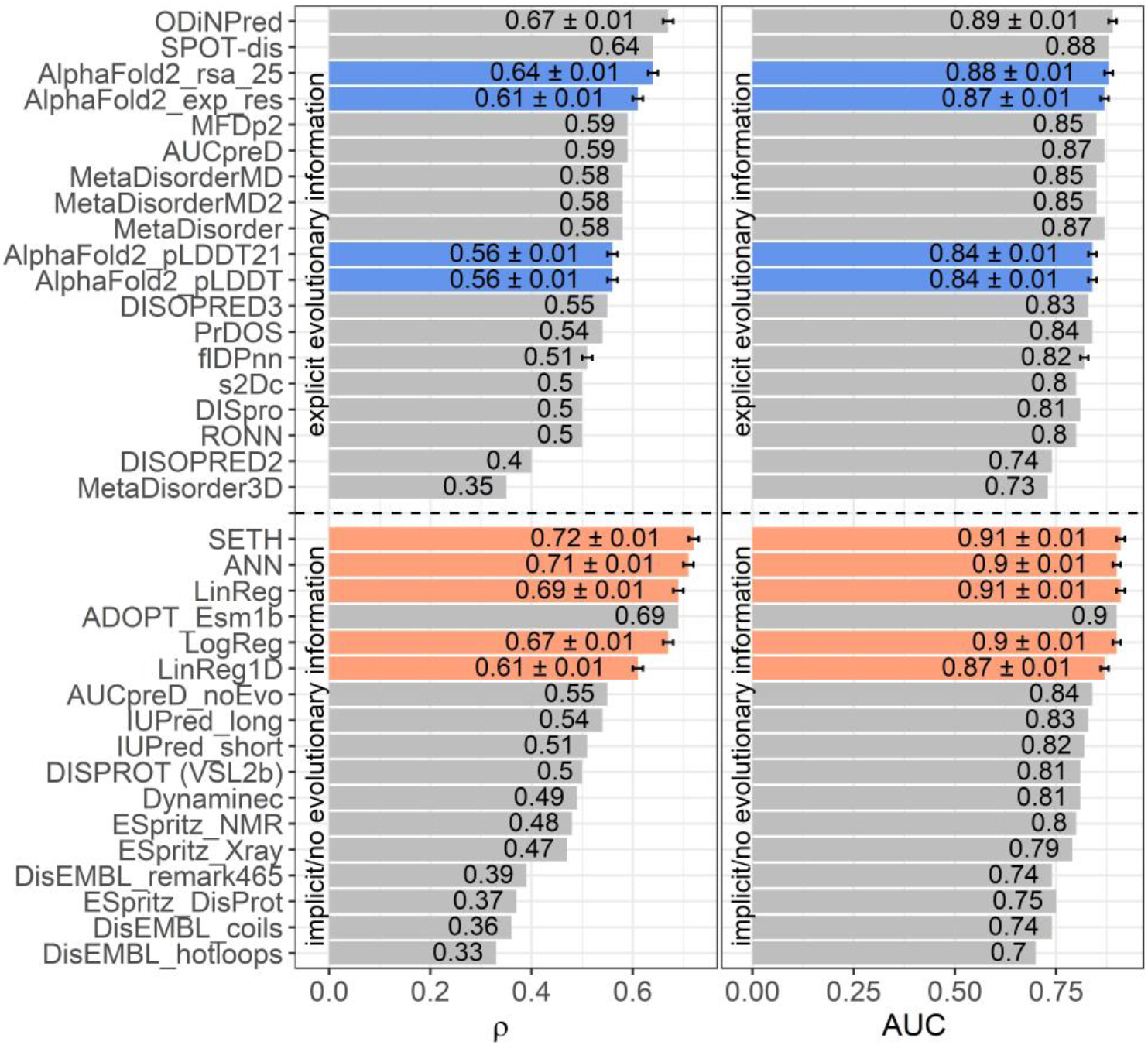
pLM-based methods performed outstandingly. Data: set CheZOD117 (Dass et al., 2020). Performances of all methods introduced here (SETH, ANN, LinReg, LogReg, LinReg1D) in orange, the ODiNPred web application in grey (Dass et al., 2020), ADOPT ESM-1b in grey (Redl et al., 2022), the fIDPnn server in grey (Hu et al., 2021) and four disorder measures derived from *AlphaFold2* (Jumper et al., 2021) in blue (these are: *AlphaFold2*_pLDDT, *AlphaFold2*_pLDDT21: smoothed over 21-consecutive residues (Akdel et al., 2021), *AlphaFold2*_exp_res: experimentally resolved prediction, *AlphaFold2*_rsa_25: running average over relative solvent accessibility averaged over 25 consecutive residues (Piovesan et al., 2022)). All other performances were taken from the previous comparison (Nielsen and Mulder, 2019) using the same test set (see Methods section 2.5). While three of our models (SETH, ANN, LinReg/LinReg1D), ADOPT ESM-1b and ODiNPred were trained on continuous chemical shift Z-scores (CheZOD scores), the logistic regression, LogReg, was trained on a binary classification of order/disorder (CheZOD score>8/≤8). ODiNPred and ADOPT ESM-1b used more proteins for training than our models. The horizontal dotted line separates models using MSAs (above line) from single sequence-based methods (below line). Error bars mark the 95% confidence interval, approximated by bootstrapping for our methods, *AlphaFold2*, the ODiNPred web application and the flDPnn server (Methods). Panel (A): Performance measured with the spearman correlation coefficient (ρ; Eqn. 2) between the ground truth and the prediction. Panel (B): Performance measured with the area under the receiver operating characteristic curve (AUC; Eqn. 3) after the binary projection of the ground truth CheZOD scores (order: CheZOD score>8, disorder: CheZOD score≤8; (Nielsen and Mulder, 2016)).

The differences between the models introduced here (LogReg, LinReg, ANN and SETH) were not statistically significant (neither for AUC nor for ρ). However, SETH had the highest mean ρ and, together with LinReg, the highest mean AUC. For a more detailed analysis, we plotted the true and predicted CheZOD scores (or for LogReg the true CheZOD scores and the predicted probability for the class “order”) for *CheZOD117* against each other in a 2D histogram for all four models (Fig. S6). SETH, ANN and LinReg agreed well with the ground truth. However, the plots revealed that SETH, LinReg and ANN tended to overestimate residue order, as indicated by the higher prediction density above the diagonal. In contrast to our other models, most of the pairs of LogReg’s predicted order probability vs. observed CheZOD scores fell into two flat clusters at 0 and 1, confirming that LogReg tended to predict extreme values optimal for classification. The removal of short disordered residues (i.e. less than 30 consecutive residues with observed CheZOD scores≤8) did not change the Spearman correlation significantly (Fig. S7).

### Shortcomings of SETH

For SETH, our best model (outperforming all others in ρ and AUC; Fig. 3), we added another analysis classifying each residue in *CheZOD117* into one of three classes according to the observed CheZOD scores: ordered (CheZOD score>8), long disorder (residues in a disorder (CheZOD score≤8) stretch with≥30 residues) and short disorder (disordered stretches with<30 residues). Firstly, SETH clearly missed short disorder (Fig. 4: predicted values for this class were approximately uniformly distributed in [0,15], with a ρ of only 0.41±0.04). Secondly, SETH overestimated order (Fig. 4 and also Fig. S6), as there was a shift of the distributions of ordered and long disordered residues to the right from the observed to the predicted scores. Thirdly, SETH predicted several residues as ordered, for which the ground truth CheZOD scores suggested long consecutive regions of disorder (Fig. 4B). For a subset of proteins, for which at least one-third of all residues were in long IDRs but SETH predicted order, *AlphaFold2*’s pLDDT largely supported our predictions of order (Fig. S8). For two of these ten proteins, we found DisProt annotations (Quaglia et al., 2022), showing disorder to order transition regions (i.e., regions that can change from disorder to order, e.g., upon binding) overlapping with the regions of wrongly predicted order (Fig. S8). Lastly, SETH predicted CheZOD scores<0 indicated long IDRs (only this class has high counts below 0, Fig. 4). This suggested zero as a second more conservative threshold for classifying disorder, to filter out short linker regions falsely labeled as disorder.

**Fig. 4:**
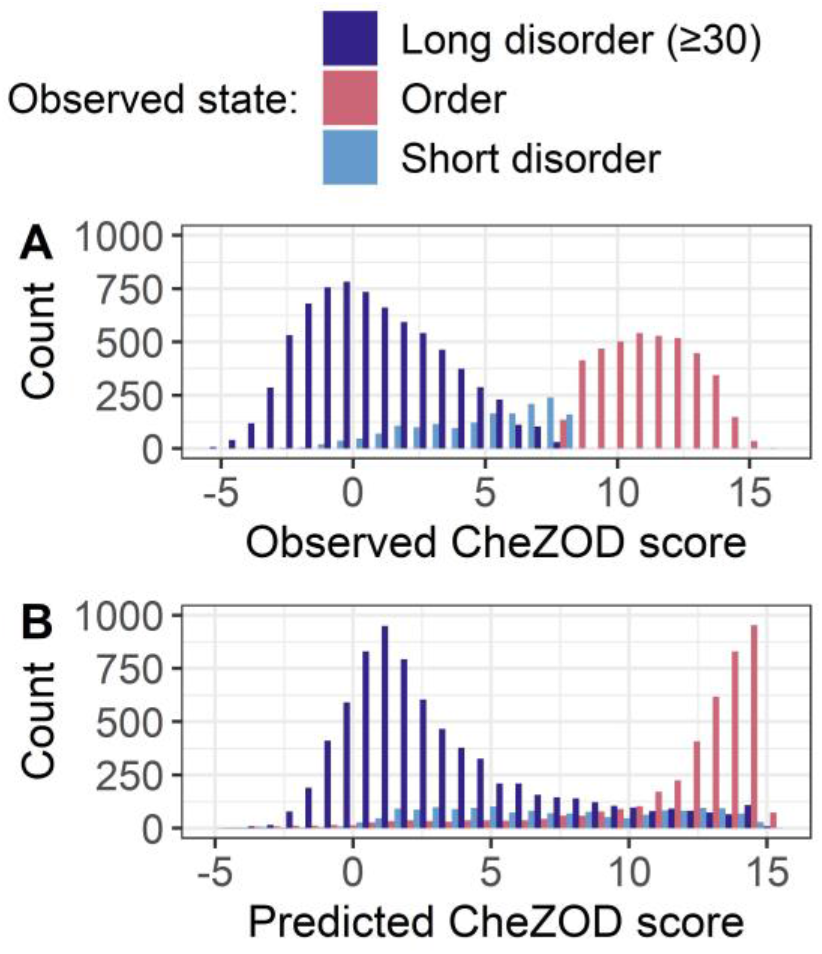
Overprediction of order and low performance for short disorder. Data: set CheZOD117 (Dass et al., 2020). All residues in CheZOD117 were classified into one of three classes: ordered (chemical shift Z-score (CheZOD score)>8), short disorder and long disorder (disordered residues (CheZOD score≤8) in a disordered region≥30), using the ground truth labels. Panel A: Distribution of the observed CheZOD scores (i.e., the ground truth labels) in the three classes. Panel B: Distribution of SETH’s predicted CheZOD scores in the three classes. SETH is a CNN trained on continuous chemical shift Z-scores and outperformed all state-of-the-art methods evaluated, as well as all our other methods.

### SETH blazingly fast

Using SETH for analyzing proteins and proteomes requires top performance (Fig. 3) and speed. On a machine with one RTX A6000 GPU with 48GB RAM, predicting the nuances of disorder for each residue of the entire human proteome (20,352 proteins) from the individual protein sequences took approximately 23 minutes. For Swiss-Prot (566,969 proteins; (The UniProt et al., 2021)), it took approximately seven hours. As a rule of thumb, SETH could predict disorder for approximately 10-20 proteins in 1 second, depending on the protein length. Even on smaller GPUs such as a single NVIDIA GeForce RTX 3060 with 12GB vRAM, computing predictions for the human proteome still took only an hour. Lastly, even on an AMD Ryzen 5 5500U CPU, performing predictions for our test set *CheZOD117* (average protein length 112) only took 12 minutes, showing that for small sets a GPU is not even necessary.

### One of 1024 embedding dimension outperformed most methods (LinReg1D)

After training, we also analyzed the regression coefficients of LinReg to better understand how ProtT5 embeddings affected the prediction. For the dimension with the highest regression coefficient (dimension 295 of 1024; Fig. S4), we subsequently plotted the raw embedding values against the true CheZOD scores (Fig. S6E) to visualize the information on order/disorder in the embeddings without supervised training. The Spearman correlation for this single dimension (ρ = 0.61) was almost the same as that for LinReg (ρ = 0.69; LinReg used all 1024 dimensions in training), showing that the pLM already learned aspects of disorder during self-supervised pre-training, i.e., without ever seeing such labels. However, in contrast to LinReg, the single dimension without supervised training avoided overestimating residue order (no accumulation of high density above the diagonal; Fig. S6).

To explicitly quantify the influence of this single most informative dimension, we additionally trained and evaluated a linear regression inputting only this 295th embedding dimension (dubbed LinReg1D). LinReg1D reached a ρ of 0.61 (LinReg ρ=0.69) and an AUC of 0.87 (LinReg AUC=0.91, Fig. 3). Therefore, this single dimension accounted for 89% or 96% of the performance of LinReg, when considering the ρ or the AUC respectively. As only a linear transformation was performed from the raw values to LinReg1D, both showed the same ρ when correlated with the true CheZOD scores.

When comparing LinReg1D to the other methods evaluated in the large-scale comparison of disorder predictors (Nielsen and Mulder, 2019), ODiNPred and ADOPT ESM-1b, even this extremely reduced model outperformed all other methods not using MSAs apart from ADOPT ESM-1b and only fell short compared to the two best-performing methods using MSAs (SPOT-Disorder (Hanson et al., 2016) and ODiNPred), when looking at both the AUC and the ρ (Fig. 3). However, compared to our other methods (SETH, LinReg, ANN, LogReg) LinReg1D performed significantly worse.

### AlphaFold2 correlated less with CheZOD scores than top methods

*AlphaFold2’s* (smoothed) predicted reliability pLDDT and its (smoothed) predicted RSA have recently been reported to capture some aspects of IDRs (Akdel et al., 2021;Wilson et al., 2021;Piovesan et al., 2022;Redl et al., 2022). However, the ρ between *AlphaFold2*’s (smoothed) pLDDT and CheZOD scores clearly neither reached the levels of the top expert solutions (SETH, LinReg, ANN, LogReg, LinReg1D, ODiNPred or ADOPT ESM-1b; Fig. 3A) trained on CheZOD scores, nor that of many other methods using MSAs (Nielsen and Mulder, 2019). Looking at the correlation between pLDDT scores and CheZOD scores in more detail (Fig. S6G) revealed that disordered residues (CheZOD score≤8) were occasionally predicted with high confidence (pLDDT>80) explaining the rather low ρ. *AlphaFold2’s* “experimentally resolved” prediction (AlphaFold2_exp_res, Fig. 4) correlated better with CheZOD scores, reaching the top 10 methods. Even better was the smoothed RSA value (ρ=0.64; AlphaFold2_rsa_25, Fig. 4), although still falling behind the top expert solutions (SETH, LinReg, ANN, LogReg, ODiNPred or ADOPT ESM-1b).

### SETH disorder predictions correlated with AlphaFold2 pLDDT

We analyzed the fitness of SETH as a fast pre-filter to distinguish between proteins/regions with low and high mean pLDDTs of *AlphaFold2* (Fig. 5). For proteins from 17 model organisms, SETH predictions correlated well with the *AlphaFold2* pLDDT (ρ = 0.67; Fig. 5A, per-organism details: Fig. S10). This trend remained after binarizing disorder using a CheZOD threshold of 8 (Fig. 5B). If the goal were to predict the classification of all proteins into those with mean pLDDT≥70 (*wanted*) and pLDDT<70 (*unwanted*), depending on the threshold in the mean predicted CheZOD score (number on the curve in Fig. 5C), this will result in different pairs of wanted proteins incorrectly missed (y-axis, Fig. 5C) given the proteins correctly ignored (x-axis, Fig. 5C). For instance, at a threshold of 8 in the mean predicted CheZOD scores, a quarter of all proteins could be avoided at an error rate of only 5% (proteins missed with pLDDT≥70). The accuracy at this threshold was 0.86. This might be relevant to prioritize/filter data in large-scale *AlphaFold2* predictions.

**Fig. 5:**
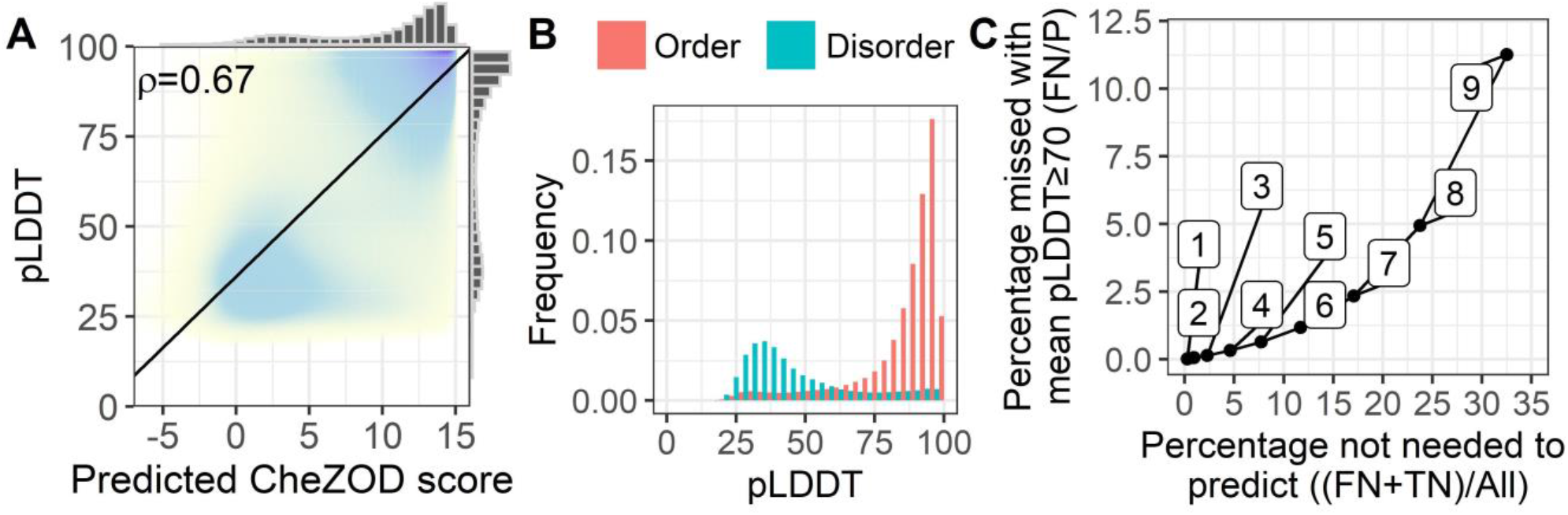
SETH’s predictions correlated with *AlphaFold2’s* pLDDT. Data: 17-ORGANISM-set (47,400,204 residues from 105,881 proteins in 17 organisms). Panel A: 2-dimensional histogram of *AlphaFold2* pLDDT (Jumper et al., 2021) against SETH disorder predictions (black line: optimal regression fit, marginal histograms on each axis; number: overall Spearman correlation coefficient ρ, Eqn. 2). Panel B: Histograms of the pLDDT of *AlphaFold2*, for the classes order (predicted CheZOD>8) and disorder (predicted CheZOD≤8). Panel C: Cost versus gain analysis using SETH as a pre-filter for *AlphaFold2*. Y-axis – Cost: The percentage of proteins with a mean predicted CheZOD score below a certain threshold (thresholds marked as numbers on the curve), but mean pLDDT≥70 (FN) out of all proteins with mean pLDDT≥70 (P). This gives the percentage of proteins with a pLDDT≥70 missed using the SETH CheZOD score prediction as a pre-filter. X-axis – Gain: The percentage of proteins with mean CheZOD score<threshold (FN+TN) out of all proteins (All). This is the percentage of proteins in the entire dataset for which *AlphaFold2* will not have to be run at all, or defines a list of priority: first run *AlphaFold2* on the proteins with lower SETH disorder. For instance, with threshold 8, a quarter of all *AlphaFold2* predictions can be avoided at an error rate of only 5%.

More importantly, the comparison of the *AlphaFold2* pLDDT and SETH’s predictions could also be used to find out more about the causes of lacking reliable *AlphaFold2* predictions. For instance, a lack of reliable *AlphaFold2* predictions was often due to disorder in proteins since low pLDDT values were mostly present for disordered residues (Fig. 5B). However, providing Fig. 5B at the organism level (Fig. S11) revealed that for some organisms, especially those with rather low mean pLDDT values (Fig. S9), SETH predicted many residues as ordered for which *AlphaFold2’s* pLDDT was low. There were even cases, where nearly the entire protein was predicted to be ordered, but *AlphaFold2* could not predict any reliable 3D structure (Fig. S12).

### Evolutionary information captured in CheZOD score distributions

Encouraged by the finding that the spectrum of predicted subcellular locations (in 10 classes) captures aspects of evolution (Marot-Lassauzaie et al., 2021b), here, we converted the CheZOD score predictions for an entire organism into a single 8-dimensional vector containing the binned normalized counts of predicted CheZOD scores. A simple PCA (Wold et al., 1987) on the resulting vectors for 37 organisms revealed a clear connection from the micro-molecular level of per-residue predictions of CheZOD-disorder to the macro-molecular level of species evolution (Fig. 6). Firstly, eukaryotes and prokaryotes (Bacteria + Archaea) were clearly separated. Secondly, even within these major groups, there appeared some relevant separation into phyla for the bacteria and into kingdoms for the eukaryotes. However, based on these limited samples, it also seemed like some groups could not be separated completely according to their disorder spectra, e.g., the fungi and the metazoa.

**Fig. 6:**
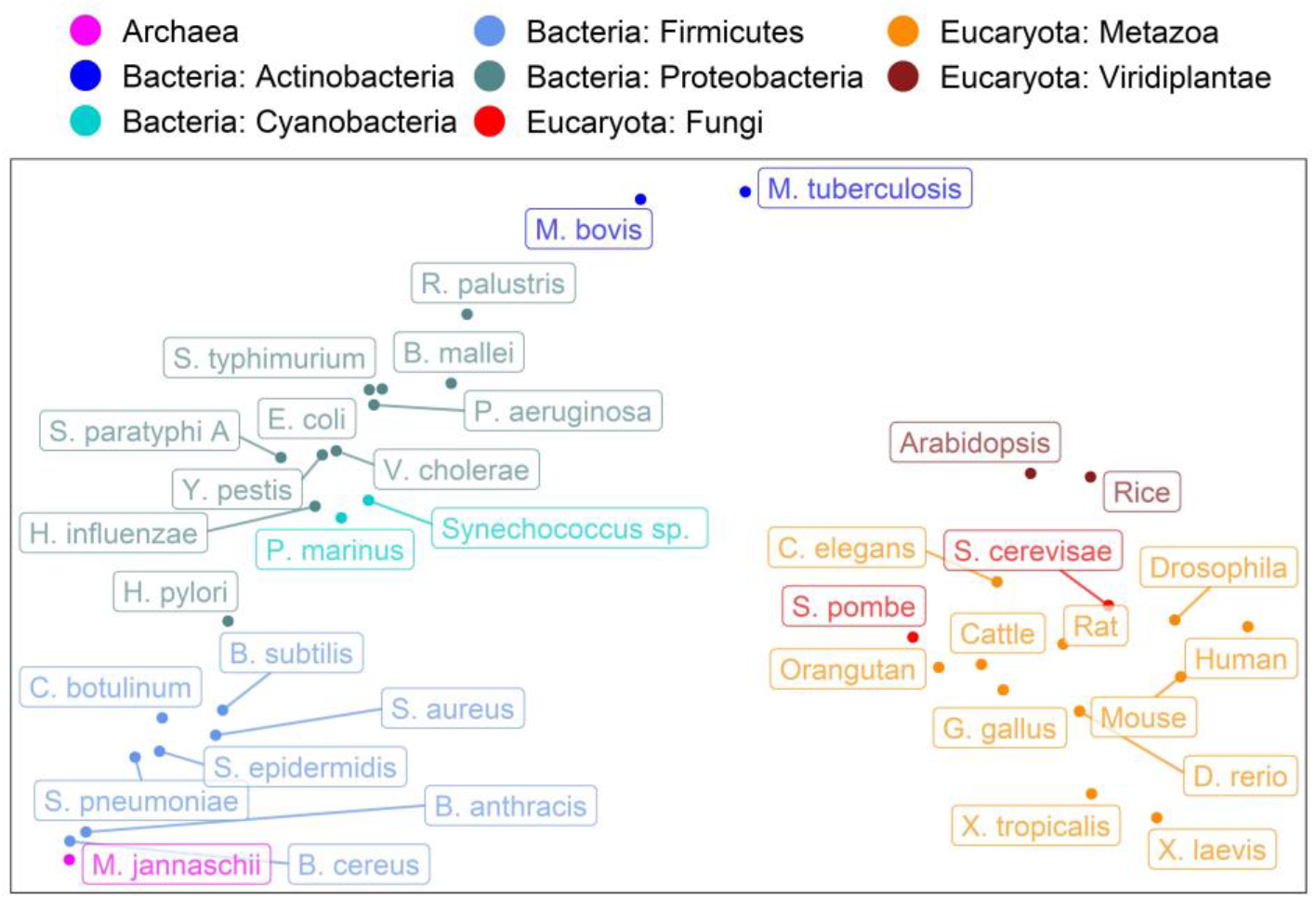
Evolution captured by spectrum of predicted CheZOD scores. Data: SETH predictions for proteins from 37 organisms taken from Swiss-Prot (The UniProt et al., 2021). The plot shows the PCA (Wold et al., 1987) of the binned spectrum of predicted CheZOD scores (8 bins, meaning 8-dimensional vectors; each vector describes one organism). The colors indicate the super-kingdoms (Eukaryota: reds, Bacteria: blues, Archaea: violet) as well as the Phyla for the Bacteria and the Kingdoms for the Eukaryotes, as given in UniProt (The UniProt et al., 2021). The organism names were shortened (for complete organism names see Table S4).

#### 4 Discussion

We introduced SETH, a shallow CNN, for predicting the continuum of residue disorder defined by CheZOD scores (i.e., the difference between observed chemical shifts from NMR and computed random coil chemical shifts (Nielsen and Mulder, 2020)). SETH’s exclusive input are embeddings from the pLM ProtT5 (Elnaggar et al., 2021). Using performance measures and data sets proposed in a recent analysis (Nielsen and Mulder, 2019), SETH outperformed three even simpler (fewer parameters) models introduced here, along with 26 other disorder prediction methods. Predictions of *AlphaFold2* have recently been shown to capture IDRs (Akdel et al., 2021;Wilson et al., 2021;Piovesan et al., 2022;Redl et al., 2022). However, we found the correlation between *AlphaFold2* predictions and CheZOD scores to be much lower than for SETH.

### Redundancy-reduction affects performance estimates, not performance

We chose our datasets (training *CheZOD1174*, and testing *CheZOD117*) and performance measures (Eqn. 2, 3) following a recent analysis (Nielsen and Mulder, 2019). However, since adequate redundancy-reduction is *sine qua non* to correctly estimate performance, we additionally removed 151 sequences from ODiNPred’s (Dass et al., 2020) training set which had, based on alignments with 80% coverage, over 20% pairwise sequence identity (PIDE) with proteins in the test set (see Methods 2.1). Unfortunately, the threshold of sequence identity T (here 20% PIDE) crucially depends on the phenotype (here disorder). In the lack of sufficiently large data sets to establish T for disorder (and many other phenotypes, including protein-protein interactions (Park and Marcotte, 2012;Hamp and Rost, 2015)), developers should be as conservative as possible. However, there is a trade-off: chose T too low, lose proteins for training/testing, chose T too high, risk substantially over-estimating performance. With our threshold, we try to balance both. However, we only removed proteins which were aligned with 80% coverage, meaning there might still be some information leakage on a smaller level (3 test proteins with PIDE>20% to training proteins at a coverage of 10%; Table S5). However, this leakage should be negligible, since none of the aligned proteins lie above the HSSP-curve (Rost, 1999). Even if there would still be some minor leakage of information, this might only balance out the over-estimates of performance of other methods, since over-estimating performance has become many times more common with the rise of AI with immense numbers of free parameters (often 10-times more parameters than samples) which can often easily zoom into residual sequence similarity between train and test set. Also considering that we used a quite conservative T, other methods tested on the same test set might more likely overestimate their performance. We cannot answer whether this over-estimate of any method is statistically insignificant or significant. That depends on many aspects of the method.

### Supervised models picked up class imbalance

The training and test sets resulting from redundancy reduction differed substantially in their distributions of CheZOD scores (Fig. S1; note the test set *CheZOD117* had not been changed, only the training set). In a binary projection, the fraction of ordered residues was 72% for the training and 31% for the testing set. Our regression models did not use any notion of classes. Thus, we could not correct for class imbalance. This might explain why our supervised regression models trained on this imbalanced data (SETH, LinReg, LinReg1D and ANN) mildly over-predicted the degree of residue order compared to the raw embedding values of dimension 295 (Fig. S6).

### Simple classification model LogReg struggled where SETH excelled

We tested the effect of increasing the model complexity when inputting only embeddings. For an ideal prediction method observed and predicted CheZOD scores would perfectly correlate, i.e., in a scatter plot with observed on the x-axis and predicted on the y-axis, perfect methods would cluster all points around the diagonal (Fig. S6). Qualitatively, our two most complex methods SETH and ANN came closest to this, followed by the simpler model LinReg, with more spread-out clusters (Fig. S6A-C). In contrast to an ideal prediction, the simplest model LogReg generated two clusters, one around probability 0 (disorder) and the other around 1 (order; Fig. S6D). Although such a bifurcation is expected for a logistic regression trained to classify, the off-diagonal shift of the data showed that LogReg struggled to capture subtle degrees of disorder/order. This qualitative analysis was supported by the ρ (Fig. 3A: SETH highest, LogReg lowest). Therefore, we established that the treatment of disorder as a regression problem (SETH, ANN, LinReg) improved over the supervised training on binary assignments (disorder/order; LogReg; Fig. 3). This was interesting because except for ODiNPred (Dass et al., 2020) and ADOPT (Redl et al., 2022), most SOTA disorder prediction methods realize a binary classification. However, the ρ was still similar between all our four models, including LogReg. Likewise, the performance on binarized CheZOD scores (order: CheZOD score>8, disorder: CheZOD score≤8), measured with the AUC did also not vary significantly. Nonetheless, SETH was consistently superior by all criteria (Fig. 3).

### Simpler, better, faster

The simplicity of a machine learning model can be proxied by the number of free parameters. Our top performing models SETH, ANN, LinReg and LogReg did not reach anywhere near the simplicity of earlier IDR prediction methods such as NORS (Liu et al., 2002) or IUPred (Dosztanyi et al., 2005) or recent adaptations of *AlphaFold2* predictions (Akdel et al., 2021;Wilson et al., 2021;Piovesan et al., 2022;Redl et al., 2022) when neglecting *AlphaFold2’s* training and only considering the disorder prediction from *AlphaFold2’s* output. Then, *AlphaFold2* binary disorder prediction would only need three parameters: choice of feature (e.g., RSA vs. pLDDT), averaging window (e.g., 25 for RSA) and a threshold (RSA<T). However, we still constrained the size of our models (Table S3). The comparison to one-hot encodings clearly demonstrated the benefit of increasing model complexity by inputting high dimensional pLM embeddings (Fig. 1). Lastly, our simplification of LinReg (LinReg1D) based on one of the 1024 dimensions of ProtT5 (Elnaggar et al., 2021), namely dimension 295 that carried 86%-96% of the signal of the entire 1024-dimensional vector (Fig. 3), reached the simplicity of very basic predictors. Still, it outperformed most complex methods.

Two of our models numerically reached higher AUC values than all other methods compared (SETH and LinReg, Fig. 3B), irrespective of whether they use MSAs or not. When considering the ρ (Fig. 3A), again two of our methods (SETH and ANN) outperformed all others. In terms of statistical significance for the ρ at the CI=95% level, all our models along with ODiNPred (Dass et al., 2020) and ADOPT ESM-1b (Redl et al., 2022) significantly outperformed all others. Of these top performers, only ODiNPred relies on MSAs, i.e., this is the only top performer for which we first need to create informative MSAs before we can analyze the disorder content of a newly sequenced proteome. Even using tools such as the blazingly fast *MMseqs2* (Steinegger and Söding, 2017), this will still slow down the analysis. In contrast, ADOPT ESM-1b also only requires pLM embeddings as input. Given the larger model used by ADOPT ESM-1b and the larger size of ESM-1b (Rives et al., 2021) compared to ProtT5 (Elnaggar et al., 2021) used by our tools, we expect the difference in speed to favor SETH more than that in performance.

### AlphaFold2 not competitive to pLM-based methods as proxy for CheZOD disorder

*AlphaFold2’s* pLDDT correlates with binary descriptions of IDRs (Akdel et al., 2021;Wilson et al., 2021;Piovesan et al., 2022). In principle, we confirmed this for CheZOD scores reflecting non-binary disorder (Fig. S6G). However, we also found *AlphaFold2* to often be certain about a predicted structure (high pLDDT) even for regions where CheZOD scores suggest long IDRs (≥30 residues; Fig. S7). One possible explanation for this might be that while *AlphaFold2* was only trained on single protein domains, some of these proteins were measured as homo- or heteromers. Consequently, the *AlphaFold2* predictions might be biased in regions that are disordered in isolation but become well-structured upon interaction. This hypothesis was supported by a very limited analysis comparing the pLDDT to DisProt annotations ((Quaglia et al., 2022); Fig. S8). Furthermore, the mean pLDDT is trivially higher for shorter than for longer proteins (Monzon et al., 2022). As proteins in the test set were shorter than average (mean sequence length in *CheZOD117*: 112), this trivial length-dependence might also explain some outliers.

Comparing several ways to utilize *AlphaFold2* predictions as a direct means to predict CheZOD scores revealed the window-averaged of the RSA to correlate even better with CheZOD scores than the prediction of “experimentally resolved” and the (smoothed) pLDDT (Fig. 3). It outperformed all but two (ODiNPred (Dass et al., 2020), SPOT-dis (Hanson et al., 2016)) of the methods not based on pLMs. However, all four methods introduced here (SETH, ANN, LinReg, LogReg) and ADOPT ESM-1b (Redl et al., 2022) topped this.

Concluding, given the many times higher runtime (we ran *AlphaFold2* (without the MSA generation step and using early stopping when one of five models reached a pLDDT≥85) and SETH on the machine with one RTX A6000 GPU with 48GB RAM and *AlphaFold2* took approximately 170 times as long as SETH), SETH appeared by far a better method for predicting disorder as defined by CheZOD scores than *AlphaFold2*. Even for the many proteins where *AlphaFold2* predictions are already available, the degree to which SETH outperformed disorder measures derived from *AlphaFold2* and the speed of SETH suggest to always use SETH instead of *AlphaFold2* to predict CheZOD-like disorder.

### Agreement between SETH’s disorder predictions and AlphaFold2’s pLDDT

*AlphaFold2*’s recent release of structure predictions (July 28, 2022), expanding the *AlphaFold2* database to over 200 million predictions, has considerably expanded the structural coverage in the protein universe. However, each day new proteins and proteomes are discovered and will require *AlphaFold2* 3D predictions. Could SETH help to prioritize how to run *AlphaFold2*, e.g., choosing the proteins most likely to have high pLDDTs (i.e., ordered proteins) first and leaving the rest for later, or completely neglecting the rest (i.e., disordered proteins)? Toward this end, we analyzed a large set of residues from 17 organisms and found the correlation between SETH’s predictions and *AlphaFold2*’s pLDDT (Fig. 5A) to be much higher than the correlation between the pLDDT and the ground truth CheZOD scores (ρ(AlphaFold2_pLDDT, ground truth)=0.56 vs. ρ(AlphaFold2_pLDDT, SETH)=0.67). This confirmed the agreement in over-prediction of order for SETH and *AlphaFold2* (Fig. S8) because if SETH and *AlphaFold2* make the same mistakes, a higher correlation is expected. These findings are at the base of using SETH to pre-filter or prioritize *AlphaFold2* predictions, e.g., using SETH protein mean CheZOD scores<8 to deprioritize or exclude some proteins will reduce costs for *AlphaFold2* by one quarter at an error rate of only 5%.

The comparisons between SETH and *AlphaFold2* also might help to rationalize some predictions, e.g., for organisms with low mean pLDDT values, SETH often predicted order where *AlphaFold2* could not predict reliable 3D structures (Fig. S9, Fig. S11). Such cases might suggest that there are some “principles of protein structure formation” not yet captured by the outstanding *AlphaFold2*. More detailed studies will have to address this speculation.

### CheZOD score disorder not equal to binary disorder

Most methods developed in the field of disorder predictions are trained on binary data: *disordered* (IDR: intrinsically disordered regions/ IDP: intrinsically disordered proteins) as opposed to *well-structured*/ *ordered*. Although this is standard procedure for machine learning, the situation for disorder is slightly different. There, we assume the set of all experimentally known 3D structures as deposited in the PDB (Burley et al., 2019) to be more representative of all well-ordered proteins than DisProt (Vucetic et al., 2005;Quaglia et al., 2022) of all disordered proteins, as the diversity of disorder is much more difficult to capture experimentally. Thus, for disorder we have many reasons to doubt that today’s experimental data are representative. This creates a “Gordian knot”: how do we train on unknown data? In previous work, we tried cutting through this knot by training on data differing from DisProt data (long loops, low contact density), but testing on DisProt (Liu et al., 2002;Schlessinger et al., 2007a;Schlessinger et al., 2007b), as, for instance, the successful method IUPred did for contacts (Dosztanyi et al., 2005). Instead, here we used CheZOD scores (Nielsen and Mulder, 2016;2019;Dass et al., 2020) introduced by Nielsen & Mulder as the “secret order in disorder”. The CheZOD perspective appealed to us because of three reasons. Firstly, it provides details or nuances for each residue. Secondly, it partially eradicates the need for a minimal threshold of continuous regions: most loops (non-regular secondary structure) of, e.g., 5-15 residues are absolutely unrelated to what we consider disorder, while loops with over 30 consecutive residues clearly fall into two distinct classes of long-loops in regular structures and disordered regions (Schlessinger et al., 2007a). Thirdly, the non-binary classification allowed to describe an entire organism by an 8-dimensional vector that captured evolution (Fig. 6).

Clearly, methods for binary disorder prediction capture information about CheZOD scores (Nielsen and Mulder, 2016;2019;Dass et al., 2020). Inversely, SETH, trained on CheZOD scores, appears to capture aspects of binary disorder (as suggested by some preliminary results from the 2nd round of CAID, the Critical Assessment of Protein Intrinsic Disorder Prediction (Necci et al., 2021)). On the other hand, one of the CAID-top methods for predicting binary disorder flDPnn (Hu et al., 2021), did not reach top rank for CheZOD scores (Fig. 3; we also queried the other CAID-top, SPOT-Disorder2 (Hanson et al., 2019) but failed to acquire results as needed). Maybe CheZOD is the “secret order in disorder”, but clearly it captures aspects somehow orthogonal to binary disorder.

### Spectra of predicted CheZOD-disorder capture rudimentary aspects of evolution

Spectra of predicted protein location capture aspects of the evolution of eukaryotes (Marot-Lassauzaie et al., 2021a). Additionally, the fraction of intrinsically disordered proteins in a proteome has been revealed as a marker for important aspects in the evolution of species (Dunker et al., 1998;Liu et al., 2002;Fuxreiter et al., 2008;Pentony and Jones, 2009;Uversky et al., 2009;Brown et al., 2011;Schlessinger et al., 2011;Vicedo et al., 2015a;Vicedo et al., 2015b). However, the single number (fraction of IDP in proteome) was too simplistic for analyses as applied to the location spectrum based on 10-dimensional vectors representing ten different subcellular compartments. The crucial step was the prediction of non-binary CheZOD scores and the idea to bin those into a spectrum with eight bins leading to 8-dimensional vectors subjected to straightforward PCA (Wold et al., 1987). Surprisingly, this already revealed a connection between the micro molecular level of per-residue CheZOD score predictions and the macro level of the evolution of species (Fig. 6). Minimally, this finding suggests that adjusting – increasing or reducing – the composition of disordered residues in proteins is a tracer of or proxy for evolutionary events. Possibly, these changes might play a role in speciation. However, at this point, the latter remains speculation. Clearly, the analysis revealed another interesting simple feature relating the micro and macro level, i.e., connecting the machinery of the proteins that shape life to the carriers of these molecular machines, namely the organisms.

## 5 Conclusions

We introduced four relatively simple novel methods exclusively using embeddings from the protein language model ProtT5 (Elnaggar et al., 2021) to predict per-residue protein disorder/order as proxied by NMR derived chemical shift Z-scores (CheZOD scores; (Nielsen and Mulder, 2020)). The best approach, dubbed SETH, captured fine-grained nuances of disorder on a continuous scale and, in our hands, appeared to outperform all compared state-of-the-art methods ((Nielsen and Mulder, 2019;Dass et al., 2020;Redl et al., 2022); Fig. 3). Our solutions were so successful because the unoptimized embeddings carried important information about disorder (Fig. 2), to the extent that mostly one of the 1024 dimensions mattered (Fig. S6E). Since SETH exclusively uses embeddings of single protein sequences, it easily scales to the analysis of entire proteomes, e.g., (dis-) order of all human proteins can be predicted in about one hour on a consumer-grade PC with one NVIDIA GeForce RTX 3060. Therefore, it enables large-scale analyses of disorder, which allowed us to show that CheZOD score distributions capture evolutionary information (Fig. 6). Although the break-through *AlphaFold2* (Jumper et al., 2021) 3D predictions are now available for most proteins, and although we could show that disorder measures of *AlphaFold2* predictions correlate with CheZOD scores, the correlation was significantly inferior to the predictions of SETH, suggesting the investment of fewer than 3 minutes per 1,000 proteins.

## 6 Software, dataset and prediction availability

SETH is available to download at https://github.com/Rostlab/SETH and available for online execution (no setup on your machine required) at https://colab.research.google.com/drive/1vDWh5YI_BPxQg0ku6CxKtSXEJ25u2wSq?usp=sharing. The predictions of SETH for Swiss-Prot (The UniProt et al., 2021) and the human proteome are available at https://doi.org/10.5281/zenodo.6673817. The training (CheZOD1174) and test dataset (CheZOD117) are also available at https://github.com/Rostlab/SETH.

## 7 Author Contributions

MH provided the AlphaFold2 structures (including pLDDT and implementation to derive the “Experimentally resolved” prediction), relative solvent accessibility values calculated from AlphaFold2 predictions, Embeddings, SETH’s predictions, and performed the MMSeqs2 clustering. MH also provided a draft of the code provided at https://github.com/DagmarIlz/SETH and at https://colab.research.google.com/drive/1vDWh5YI_BPxQg0ku6CxKtSXEJ25u2wSq?usp=sharing, which was refined by DI. DI trained and tested all models and analyzed the data with help of MH. DI wrote the first manuscript draft and generated the figures; iteratively, MH, DI and BR refined the manuscript. BR helped in the study planning, design and setting the focus. All authors read and approved the final manuscript.

## 8 Funding

This work was supported by the Bavarian Ministry of Education through funding to the TUM and by a grant from the Alexander von Humboldt foundation through the German Ministry for Research and Education (BMBF: Bundesministerium für Bildung und Forschung), by two grants from BMBF (031L0168 and program “Software Campus 2.0 (TUM) 2.0” 01IS17049) as well as by a grant from Deutsche Forschungsgemeinschaft (DFG-GZ: RO1320/4-1).

## 9 Acknowledgments

Particular thanks to Keith Dunker (Indiana U) for his help in realizing the importance of the topic of IDRs and IDPs. Further thanks primarily to Tim Karl (TUM) for invaluable help with hardware and software and to Inga Weise (TUM) for crucial support with many other aspects of this work. Thanks to Maria Littmann and Michael Bernhofer (both TUM) for listening, helping and contributing their insights in discussions and thank you to Konstantin Schütze (TUM) for his input in the review process. Last, but not least, thanks to all those who contribute toward maintaining public databases, and to all experimentalists who enabled this analysis by making their data publicly available.

## 10 Abbreviations

3D: three-dimensional, i.e., coordinates of all atoms/residues in a protein
AI: artificial intelligence
*AlphaFold2*: AI-based method reliably predicting protein 3D structure from MSAs (Jumper et al., 2021)
ANN: an artificial feed-forward neural network
AUC: area under the receiver operating characteristic curve
CheZOD scores: chemical shift Z-scores from NMR (Nielsen and Mulder, 2019)
CI: confidence interval, here typically used as the 95% CI implying an interval between ±1.96*Standard Error
CNN: convolutional neural network
*ColabFold*: protocol for fast execution of *AlphaFold2* (Mirdita et al., 2022)
GPU: graphical processing unit
IDP: intrinsically disordered proteins (Dunker et al., 2013)
IDR: intrinsically disordered regions (Dunker et al., 2013)
LinReg: a linear regression model
LogReg: a logistic regression model
MSA: multiple sequence alignment
NMR: nuclear magnetic resonance
PCA: principle component analysis
PIDE: percentage pairwise sequence identity
pLDDT: predicted local distance difference test from *AlphaFold2* (Jumper et al., 2021)
pLM: protein Language Model
ProtT5: particular pLM (Elnaggar et al., 2021)
RSA: relative solvent accessible surface area of a residue
SETH: a CNN for continuous disorder prediction (our best model)
SOTA: state-of-the-art
t-SNE: t-distributed stochastic neighbor embedding
ρ: Spearman correlation coefficient.

## Notes

https://github.com/DagmarIlz/SETH

https://doi.org/10.5281/zenodo.6673817

### Competing Interest Statement

The authors have declared no competing interest.

### Summary of Updates

Major changes: (1) Additional baseline: one-hot encodings (2) Additional analysis and discussion of redundancy reduction (3) Addition of flDPnn in evaluation (4) Added discussion of CheZOD scores against binary disorder

