## Supplementary Material for "SETH predicts nuances of residue disorder from protein embeddings"

### 1 Short description of the Supplementary Material

This Supplementary Material extends our work on creating a predictor for nuances of protein residue disorder, as defined by chemical shift Z-scores (CheZOD scores; (Nielsen and Mulder, 2019)), using only embeddings of single protein sequences derived from protein language models as input. Firstly, we describe the implementation and optimization of our methods not performing best (1.1 Supplementary Methods). Secondly, we analysed the distribution of the labels (the CheZOD scores) in the training (CheZOD1174; Fig. S1A) and test dataset (CheZOD117; Fig. S1B) used throughout our analyses. Table S5 shows some residual similarities between these two sets. Further, we provide results for the hyperparameter optimization of the neural network (dubbed ANN; Fig. S2) and the convolutional neural network (dubbed SETH; Fig. S3) trained in our work. Table S3 provides the resulting number of parameters for all our models. We also show an in-depth analysis of the linear regression model (dubbed LinReg) that we trained on ProtT5 embeddings (Elnaggar et al., 2021), where we analysed the most important embedding dimensions defined by the largest regression coefficients and the Spearman correlation coefficients between the true CheZOD scores and single dimensions of the ProtT5 embeddings for the residues in the test set (Fig. S4). Furthermore, we show the full names of abbreviated organisms of Fig. 6 (Table S4), where we analysed SETH's predictions on capturing evolution.

In addition, we expand our evaluation by providing a visualization of the receiver operating characteristic curves for the evaluated models (ROC curves; Fig. S5) and by providing a detailed plot of the correlation of various predictions with the CheZOD scores (Fig. S6). Also, we trained and evaluated baseline models on random embedding vectors sampled from a normal distribution for all our model types (Table S1). As a last additional evaluation step, we checked if the performance of the evaluated models changed when excluding short disordered residues (disordered region<30; Fig. S7) and added an analysis on proteins in our test set, where SETH predicted order, but according to the ground truth values the residues were in a long disordered region (disordered region≥30; Fig. S8).

Expanding our analyses on AlphaFold2's pLDDT (Jumper et al., 2021), firstly, we provide a table with information on 19 organisms for which we downloaded Alphafold2 structures (Table S2). Secondly, we show mean pLDDT values for the 19 organisms (Fig. S9) and for 17 out of these 19 organisms we additionally display the agreement between the pLDDT and SETH's predicted CheZOD scores, both on a continuous scale and binned into the classes order and disorder based on SETH's predicted CheZOD scores (Fig. S10, S11). Lastly, analysing proteins which did not show an agreement between the pLDDT and SETH's predictions in more detail, Fig. S12 shows the CheZOD score distribution and AlphaFold2 prediction for a protein where AlphaFold2 could not predict anything with high reliability, but SETH predicted order.

### 1.1 Supplementary Methods

**Implementation and optimization of disorder predictors.** This section expands on methods section 2.3 of our main text. LinReg, ANN and LogReg, three of our models, were implemented with scikit-learn (Pedregosa et al., 2011). For LinReg the *LinearRegression* model was used. For ANN, the *MLPRegressor* model was used and lastly, for LogReg, the *LogisticRegression* model was used. For the linear regression, all parameters were left at default levels. For LogReg, the class imbalance between ordered/disordered residues was taken into account by setting "class_weight" to balanced, which causes weighting inversely proportional to class frequency. Furthermore, the maximal number of iterations was chosen to be 400, while all other parameters were left at standard values. For ANN, the tanh was used as the activation function between hidden layers, an identity function was used in the final output layer, the random state was set to 1 to guarantee reproducibility and early stopping was activated. Furthermore, an optimization of the following hyperparameters was performed using scikit-learn's (Pedregosa et al., 2011) GridSearchCV on CheZOD1174: a) number of hidden neurons, b) solver (adam (Kingma and Ba, 2017) versus stochastic gradient descent (SGD) (Rumelhart et al., 1986)), and c) the learning rate. The ρ between the true and predicted CheZOD scores was used as the scoring function in this optimization. The best performing model (in terms of mean ρ), was used for any further analysis, resulting in the stochastic gradient descent, 2 hidden layers with 3 neurons each and a constant learning rate of 0.001 (Fig. S2).

### 1.2 Supplementary Figures

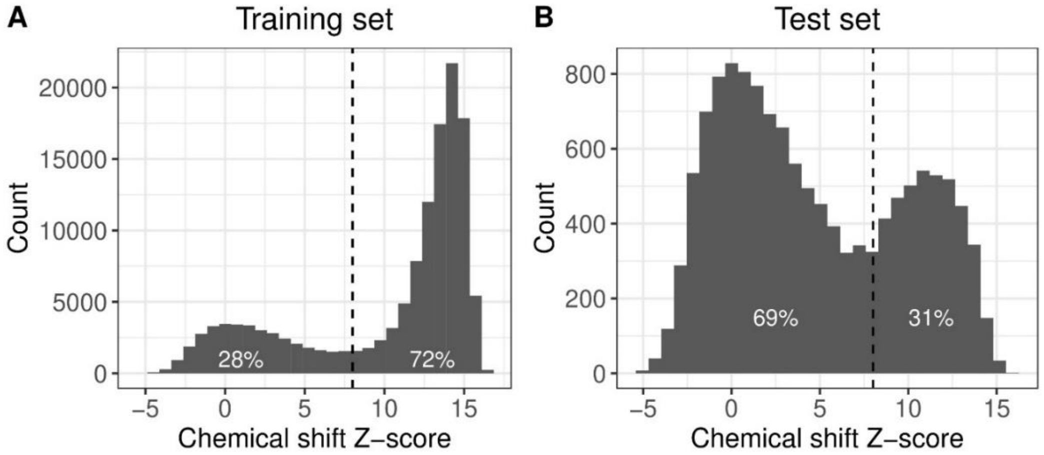

**Fig. S1: Label distribution in training and test set.** Distribution of the chemical shift Z-scores (CheZOD scores) in (A) our training set CheZOD1174, which contains 1174 sequences and 132,545 residues and (B) our test set CheZOD117, which contains 117 sequences and 13,069 residues. The dotted line separates ordered (CheZOD score > 8; 72% for CheZOD1174, 31% for CheZOD117) from disordered residues (CheZOD score ≤ 8; (Nielsen and Mulder, 2016)). The data originates from (Dass et al., 2020).

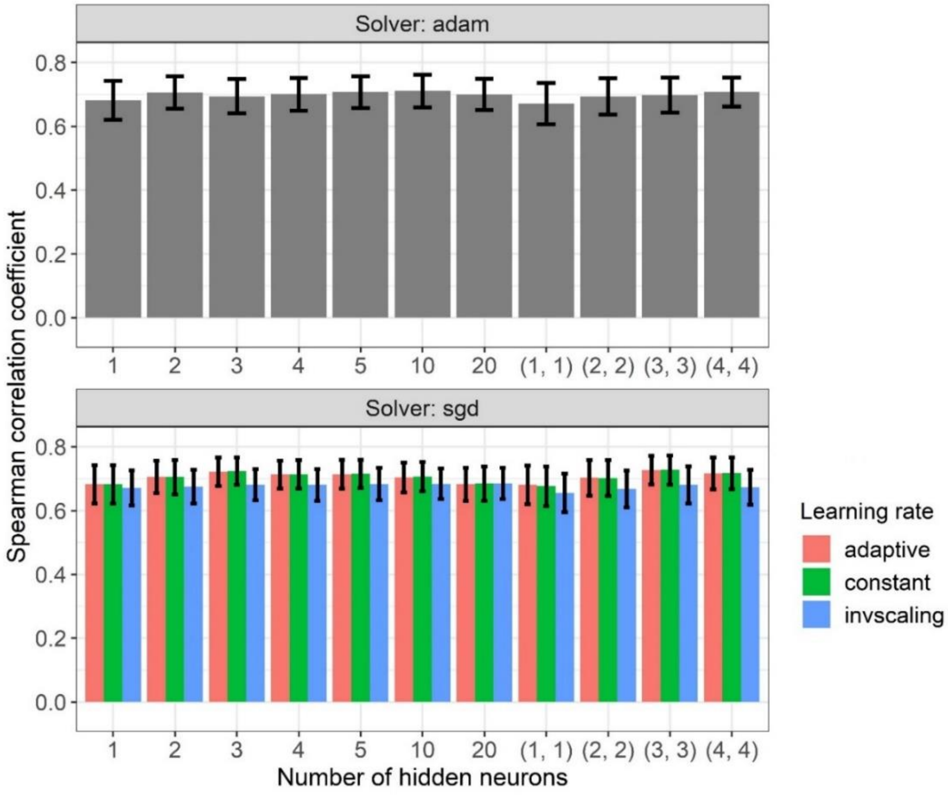

**Fig. S2: Hyperparameter optimization ANN.** Data: CheZOD1174 (Dass et al., 2020). Shown are the hyperparameter optimization results of the neural network ANN. The optimization was performed with GridSearchCV of scikit-learn. The Spearman correlation coefficient between the true and predicted chemical shift Z-scores was used as the scoring function in this optimization. The error bars mark the standard error as reported through GridSearchCV in the variable "std_test_score".

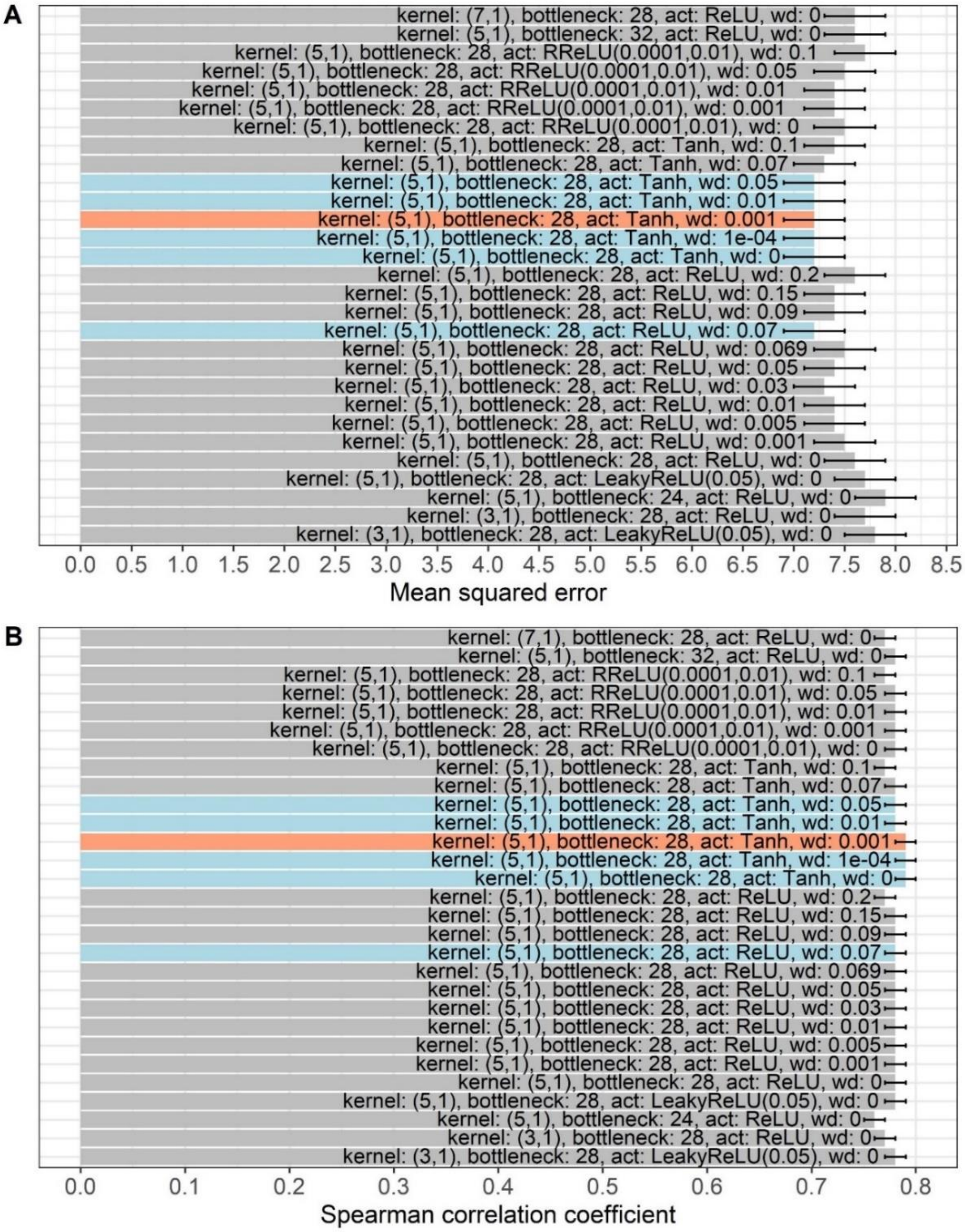

**Fig. S3: Hyperparameter optimization SETH.** Data: CheZOD1174 (Dass et al., 2020). Visualized are the hyperparameter optimization results of the convolutional neural network SETH. All models were trained on 90% of the proteins of CheZOD1174 (Dass et al., 2020) that were randomly chosen and validated on the remaining 10%. The (B) Spearman correlation coefficient between the true and predicted chemical shift Z-scores and the (A) mean squared error between the true and predicted chemical shift Z-scores, which was used as the loss function in the optimization, were used to decide on the final model. The errors mark the 95% confidence intervals approximated by multiplying 1.96 with the bootstrap standard deviation. In blue, the models performing best in terms of the mean squared error are marked. The final model is marked in red. The variables varied in the optimization were the kernel size (dubbed kernel), the number of output channels of the first convolutional layer (dubbed bottleneck), the activation function between the two convolutional layers (dubbed act) and the weight decay parameter in the optimizer (dubbed wd).

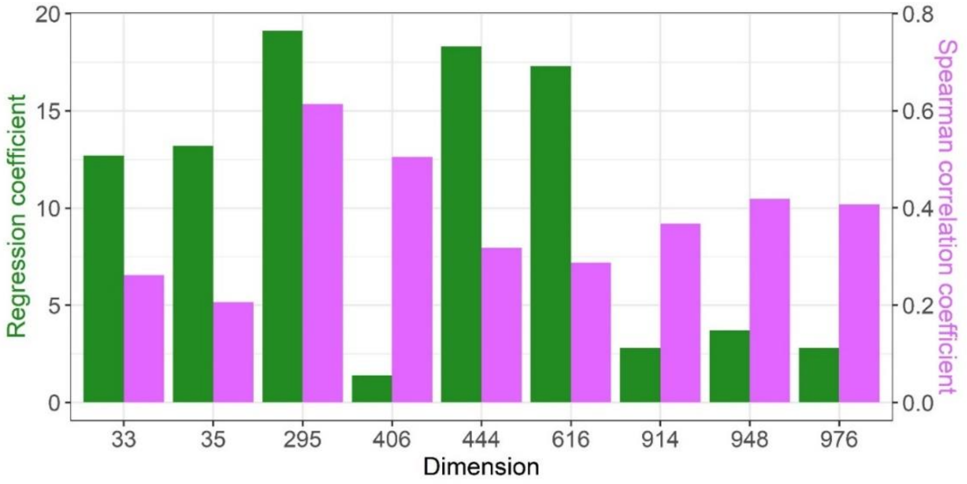

**Fig. S4: Embedding dimension 295 most important.** Absolute value of the regression coefficients of LinReg and the Spearman correlation coefficients between the true chemical shift Z-scores and the values of a selected dimension of the ProtT5 embeddings (Elnaggar et al., 2021) for the residues in CheZOD117 (Dass et al., 2020). Shown are these values for the dimensions which have one of the five largest Spearman correlation coefficients and/or one of the five largest regression coefficients. The regression coefficient and the Spearman correlation coefficient indicate different dimensions as important in predicting disorder (apart from dimension 295).

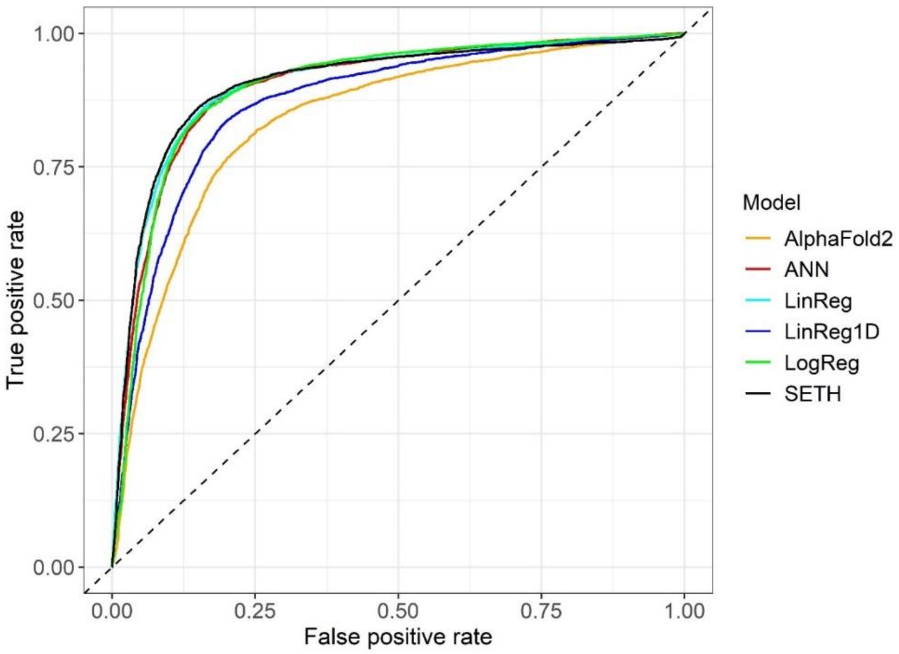

**Fig. S5: ROC curves.** Data: set CheZOD117 (Dass et al., 2020). Receiver operating characteristic (ROC) curve for the disorder predictors created here and AlphaFold2's pLDDT (Jumper et al., 2021). SETH, LinReg, LinReg1D and ANN are regression models, while LogReg is a classification model. To receive the ROC curves, the continuous chemical shift Z-scores were mapped to the classes order (chemical shift Z-score > 8) and disorder (chemical shift Z-score ≤ 8). The ROC curves show no abnormalities.

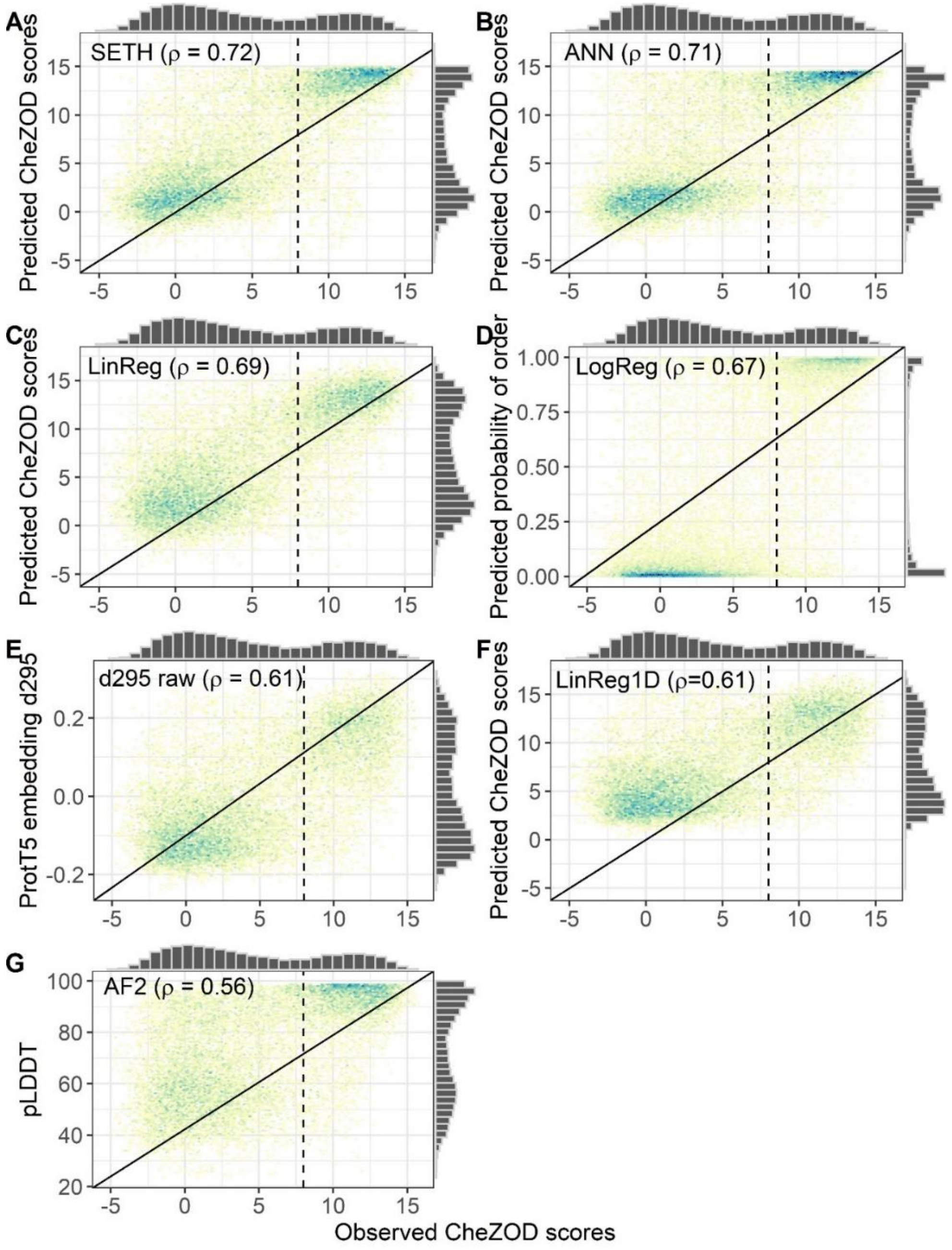

**Fig. S6: Correlation of predictors with ground truth.** Data: set CheZOD117 (13,069 residues; (Dass et al., 2020)). 2-dimensional histogram of the observed chemical shift Z-score (CheZOD score; (x-axis)) against predictions for seven methods (y-axis). Additionally, a marginal histogram is visible for each axis. Methods introduced here: (A) SETH, (B) ANN, (C) LinReg, (D) LogReg, (E) raw embedding dimension 295 (d295 raw) of ProtT5, which is the dimension with the highest regression coefficient in LinReg and (F) LinReg1D, a linear regression trained on the most important embedding dimension (d295 raw) for disorder prediction, based on the regression coefficients of LinReg. Additionally, Panel (G) shows AlphaFold2 (AF2; (Jumper et al., 2021)). The black diagonal in each plot marks the optimal regression fit. Vertical dotted black lines separate ordered (CheZOD score>8) from disordered residues (CheZOD score≤8; (Nielsen and Mulder, 2016)). The Spearman correlation coefficient (ρ) was estimated from bootstrapping in all panels. For all panels except panel (D), the same colors correspond to the same number of points in an area. The plot shows that nuances of per-residue CheZOD scores are accurately predicted by our models.

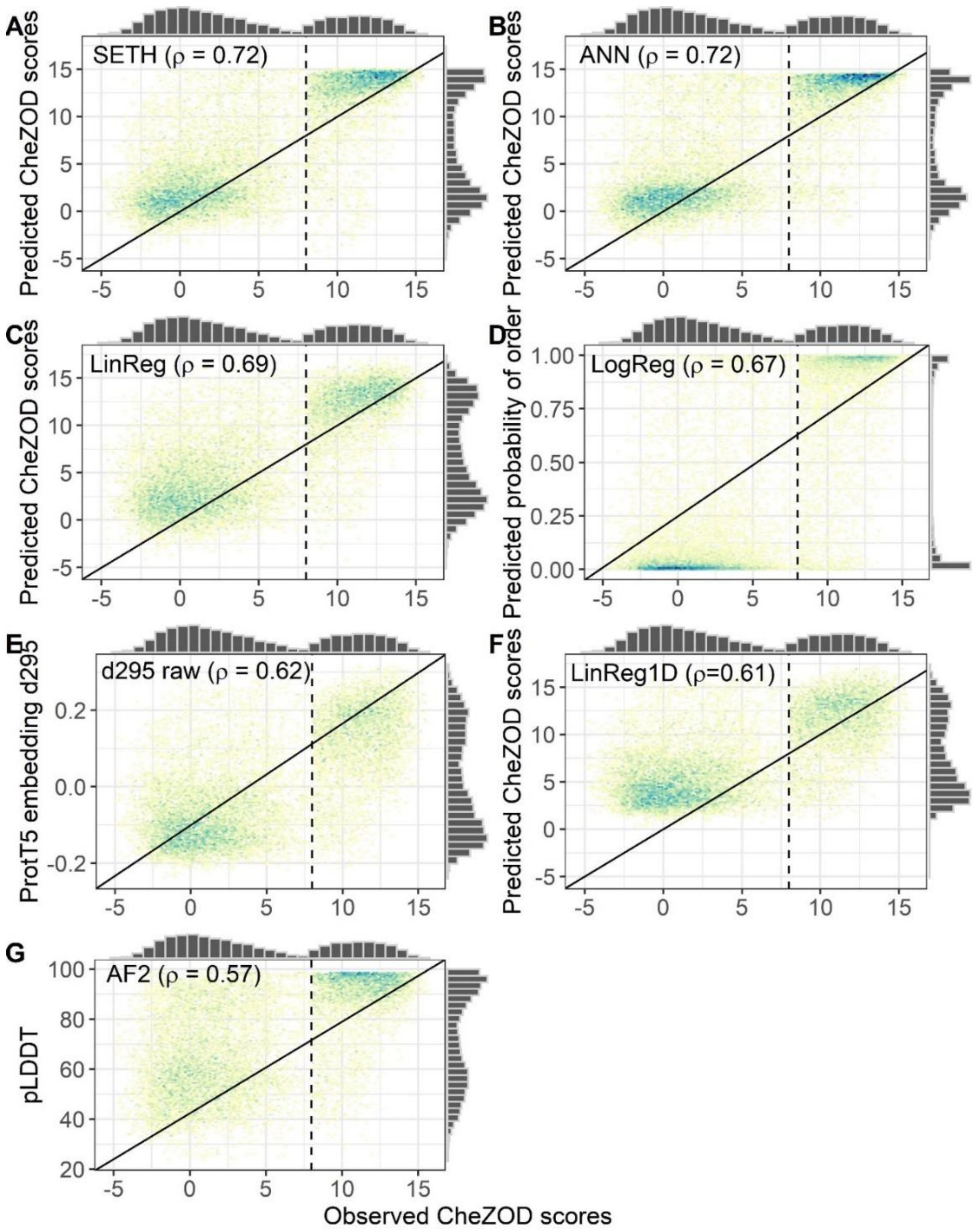

**Fig. S7: Correlation of predictors with ground truth, excluding short disorder.** Data: set CheZOD117 (13,069 residues; (Dass et al., 2020)), without short disordered residues (short disordered residues: disordered regions<30; disorder: Chemical shift Z-score (CheZOD score≤8); resulting dataset: 11414 residues). 2-dimensional histogram of the observed CheZOD score (x-axis) against predictions for seven methods (y-axis). Additionally, a marginal histogram is visible for each axis. Methods introduced here: (A) SETH, (B) ANN, (C) LinReg, (D) LogReg, (E) raw embedding dimension 295 (d295 raw) of ProtT5, which is the dimension with the highest regression coefficient in LinReg and (F) LinReg1D, a linear regression trained on the most important embedding dimension for disorder prediction (d295 raw), based on the regression coefficients of LinReg. Additionally, Panel (G) shows AlphaFold2 (AF2; (Jumper et al., 2021)). The black diagonal in each plot marks the optimal regression fit. Vertical dotted black lines separate ordered (CheZOD score>8) from disordered residues (CheZOD score≤8; (Nielsen and Mulder, 2016)). The Spearman correlation coefficient (ρ) was estimated from bootstrapping in all panels (95% Confidence interval is ρ ±0.01 for all panels). For all panels except panel (D), the same colors correspond to the same number of points in an area. The plot shows that excluding the short disordered residues does not significantly change the performance (compare to Fig.

S6).

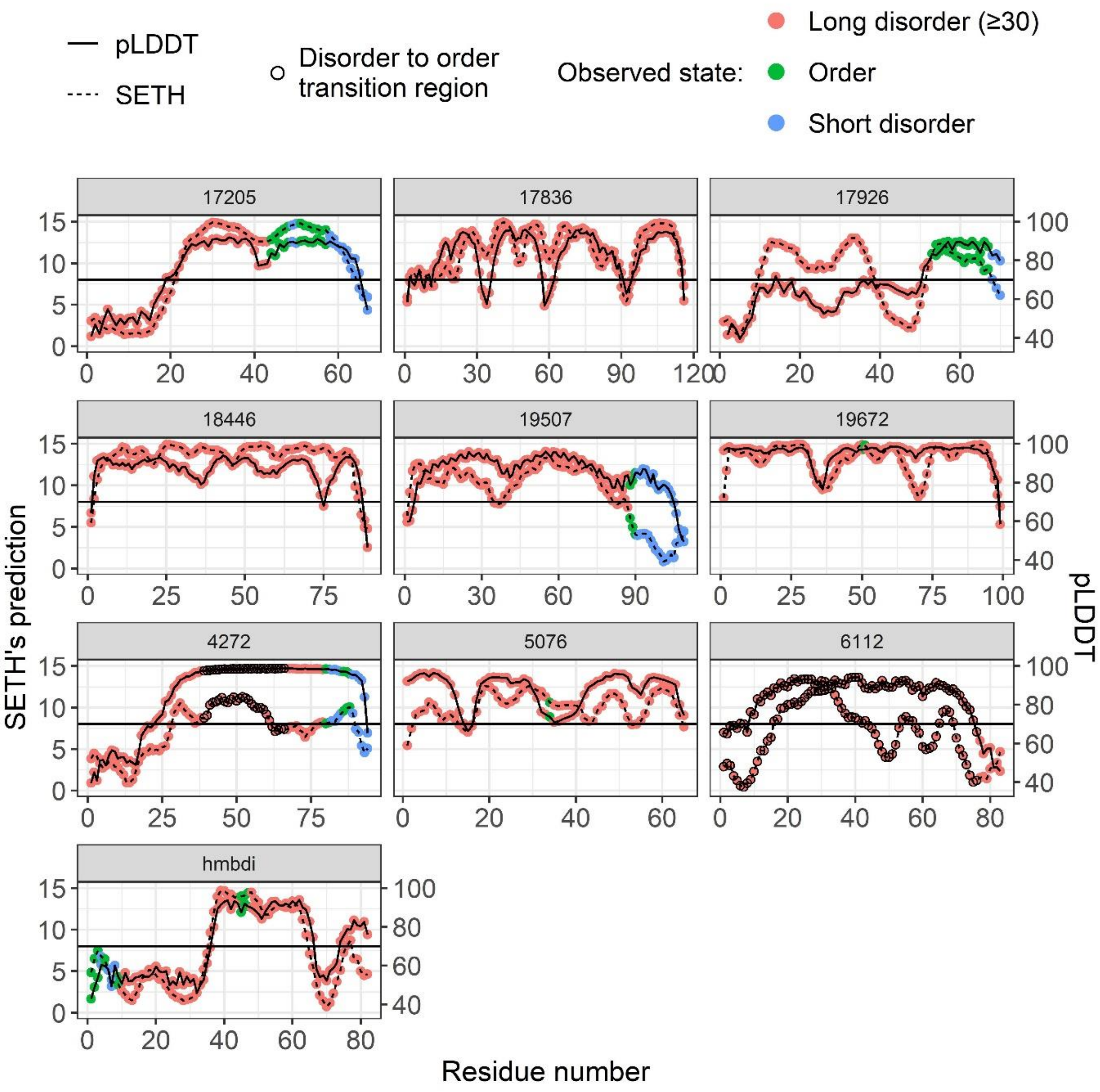

**Fig. S8: Overprediction of order.** Data: proteins from CheZOD117 (Dass et al., 2020), where at least one-third of residues fulfill the following criteria: predicted chemical shift Z-scores (CheZOD scores) hint at ordered (CheZOD>8) but according to ground truth it is a long disorder stretch (CheZOD score≤8 in a region≥30 residues). For each protein, SETH's predicted CheZOD scores are plotted along the sequence (dotted line), together with AlphaFold2's pLDDT (solid line; (Jumper et al., 2021)). The colors show the class of the residue according to the ground truth labels. The classes are: order (CheZOD score>8), short and long disorder (CheZOD score≤8) in a region of less than or more than 30 residues, respectively. The circles in the plots for proteins 4272 and 6112 mark the DisProt annotation (Quaglia et al., 2022) disorder to order transition. The horizontal black line marks the threshold between disorder and order in the CheZOD scores (order: CheZOD score>8; order: CheZOD score ≤8; (Nielsen and Mulder, 2016)) and the threshold between confident and unreliable predictions (threshold: 70) in the pLDDT.

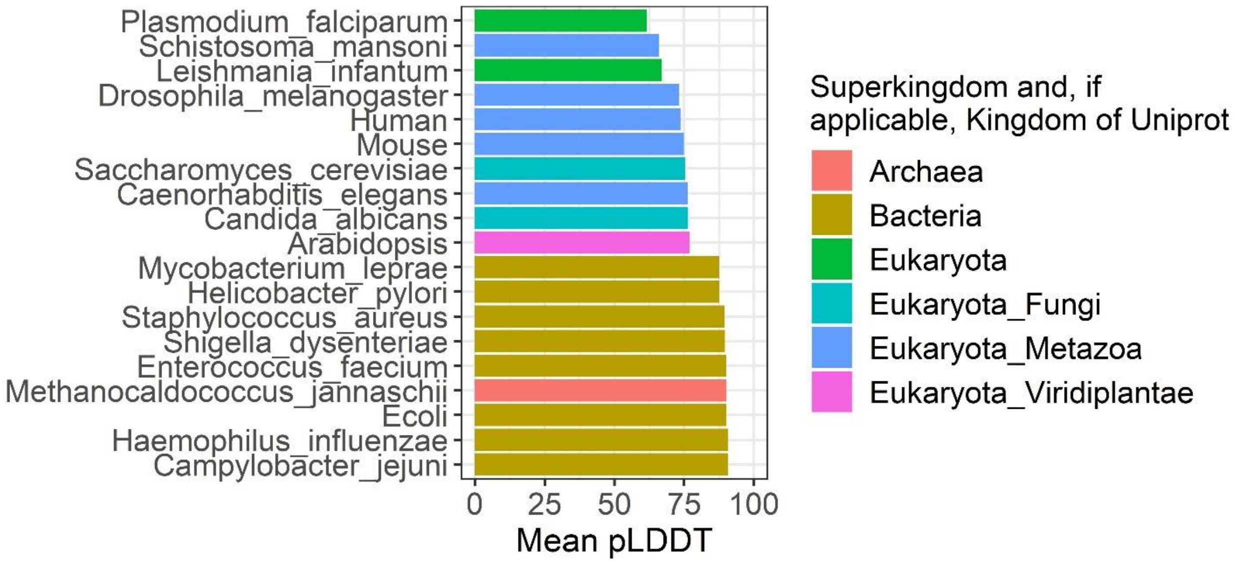

**Fig. S9: Mean pLDDTs of 19 organisms.** Data: All residues in the available proteins with extractable pLDDT values (for some proteins the table format was inconsistent and therefore, pLDDT values could not be extracted. These were excluded.) from the 19 organisms downloaded from the AlphaFold2 database (Table S2; (Jumper et al., 2021)), as described in Methods section 2.6 - Comparison: CheZOD score predictions and pLDDT in 17 organisms. Displayed are the mean pLDDT values per organism, calculated over all residue pLDDT values of the organism. The color shows the Superkingdom and if the organism had an annotated Kingdom, the Kingdom of UniProt (The UniProt et al., 2021). Some organism names were shortened in the plot: Human=*Homo sapiens*, Mouse=*Mus musculus*, Ecoli=*Escherichia coli*, Arabidopsis=*Arabidopsis thaliana*.

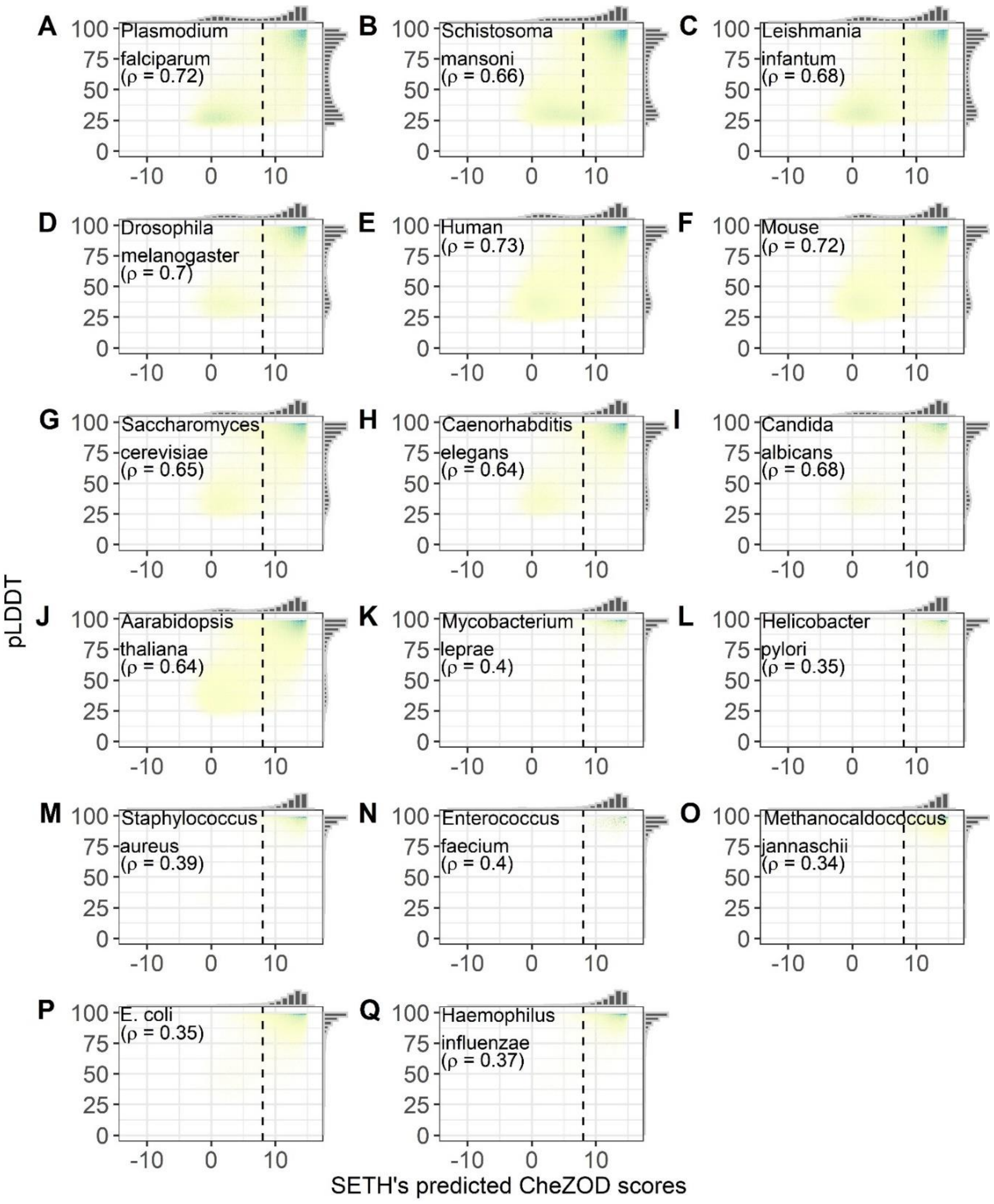

**Fig. S10: Correlation of SETH and AlphaFold2 for 17 organisms.** Data: 17-ORGANISM-set (Methods section 2.6 - Comparison: CheZOD score predictions and pLDDT in 17 organisms; 105,881 proteins and 47,400,204 residues from 17 organisms; Table S2). 2-dimensional histogram of the pLDDT of AlphaFold2 (Jumper et al., 2021) versus disorder predictions of SETH. Additionally, a marginal histogram is visible for each axis. The Spearman correlation coefficient (ρ) displayed was calculated with Eqn. 2 (Methods section 2.5). Vertical dotted black lines separate predicted ordered (CheZOD score>8) from predicted disordered residues (CheZOD score≤8; (Nielsen and Mulder, 2016)). Some organism names were shortened in the plot: Human=*Homo sapiens*, Mouse=*Mus musculus*, E. coli=*Escherichia coli*.

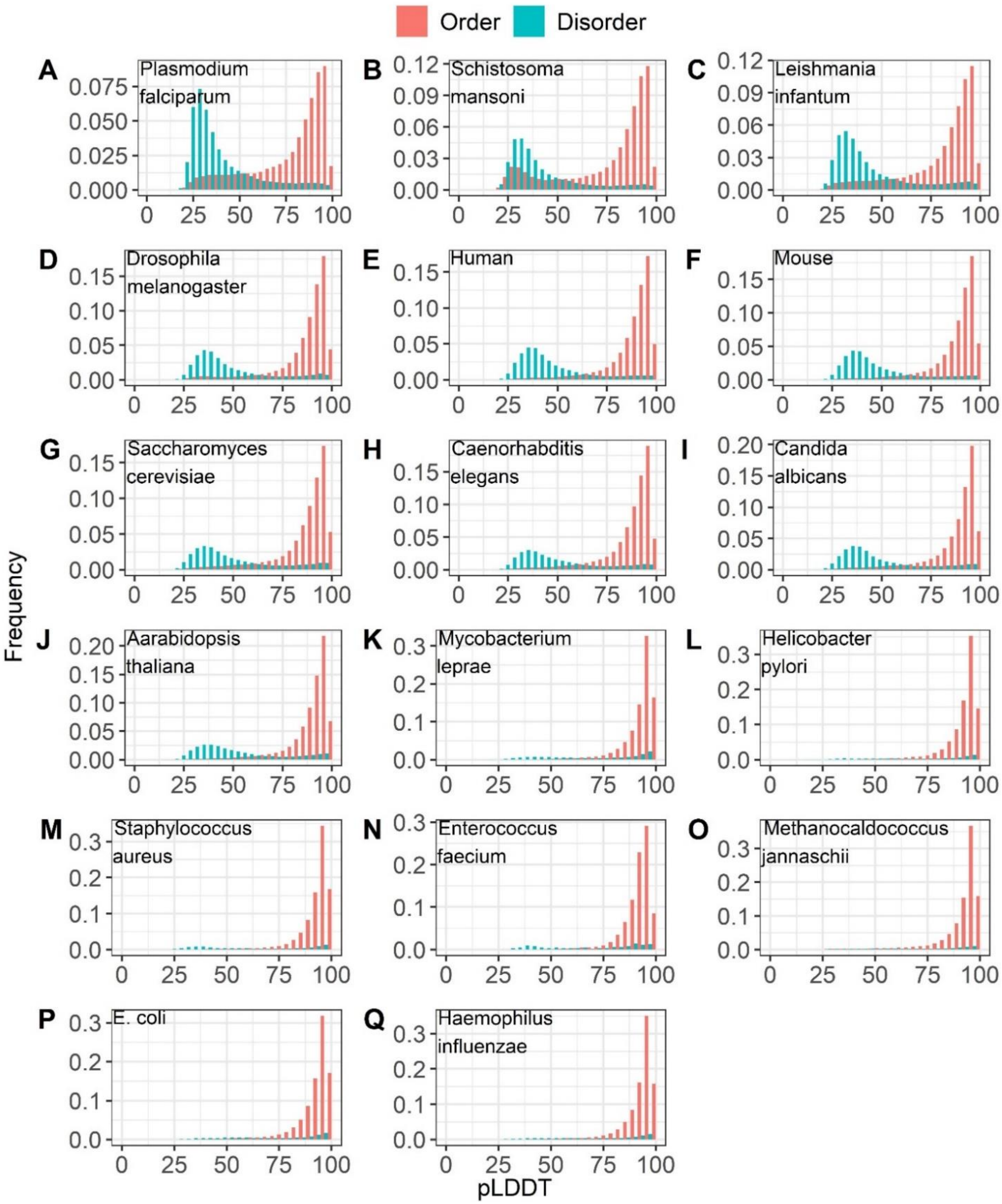

**Fig. S11: Binarized agreement between SETH and AlphaFold2 for 17 organisms.** Data: 17-ORGANISM-set (Methods section 2.6 - Comparison: CheZOD score predictions and pLDDT in 17 organisms; 105,881 proteins and 47,400,204 residues from 17 organisms; Table S2). Histograms of the pLDDT of AlphaFold2, for the classes order and disorder, classified by using a threshold of 8 in the predicted CheZOD scores of SETH (order: CheZOD score>8, disorder: CheZOD score≤8; (Nielsen and Mulder, 2016)). Some organism names were shortened in the plot: Human=*Homo sapiens*, Mouse=*Mus musculus*, E. coli=*Escherichia coli*.

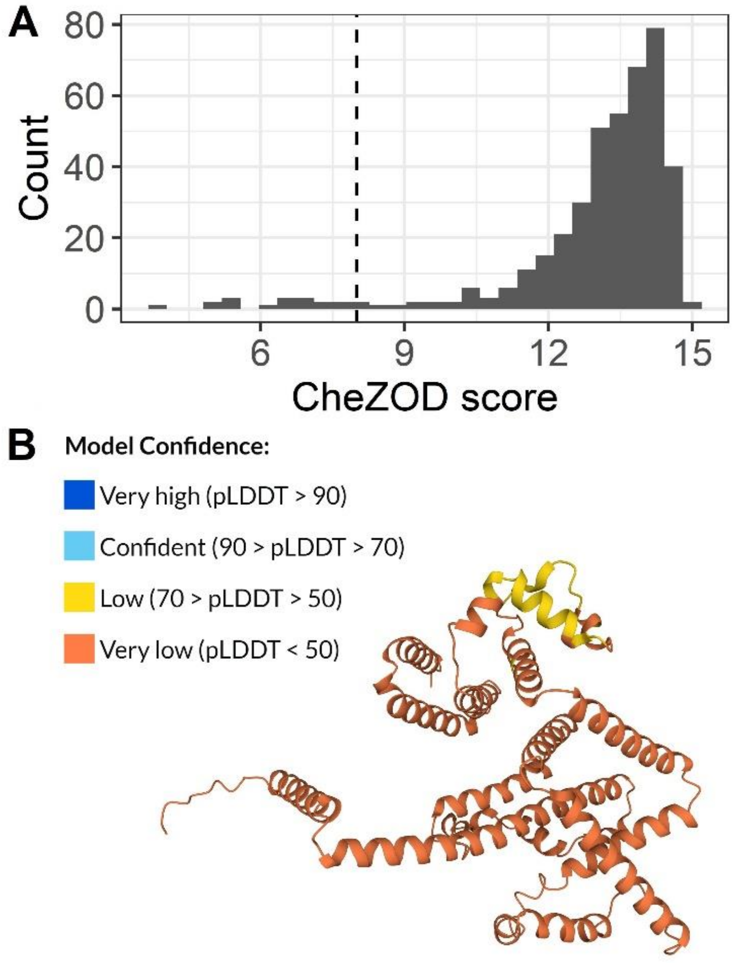

**Fig. S12: Exceptional disagreement between SETH and AlphaFold2.** Data: Protein with UniProt ID Q8ILI5 of Plasmodium falciparum. Panel A: Histogram of SETH's predicted chemical shift Z- scores (CheZOD scores) for the protein. The vertical dotted black line separates predicted ordered (CheZOD score>8) from predicted disordered residues (CheZOD score≤8; (Nielsen and Mulder, 2016)). Panel B: AlphaFold2 (Jumper et al., 2021) predicted structure of the protein. On the grander scale, proteins which were predicted to be ordered, but AlphaFold2 could not predict a reliable structure were rare since there is a general agreement between the pLDDT and SETH's disorder predictions (Fig. 5A).

### 1.3 Supplementary Tables

**Table S1: Baseline model performance.** Shown are the performances of the random baseline models for the models created here, established by replacing CheZOD1174 in training and CheZOD117 in testing by datasets of the same size but consisting of random embeddings of shape 1024 sampled from a standard normal distribution. The performance is measured by the Spearman correlation coefficient (ρ) and the area under the receiver operating characteristic curve (AUC). The errors mark the 95% confidence intervals approximated by multiplying 1.96 with the bootstrap standard deviation. To calculate the AUC, the continuous chemical shift Z-scores (CheZOD scores) were mapped to the classes order (CheZOD score > 8) and disorder (CheZOD score ≤ 8). As expected, the ρ is never significantly different from zero and the AUCs are not significantly different from 0.5.

| Baseline model | ρ | AUC |
|---|---|---|
| SETH | -0.02 ± 0.02 | 0.49 ± 0.01 |
| LinReg | -0.01 ± 0.02 | 0.49 ± 0.01 |
| LinReg1D | -0.003 ± 0.02 | 0.50 ± 0.01 |
| LogReg | -0.001 ± 0.02 | 0.50 ± 0.01 |
| ANN | -0.02 ± 0.02 | 0.50 ± 0.01 |

**Table S2: Organism information for Alphafold2 download.** Information for predictions of 19 organisms downloaded from AlphaFold2 (Jumper et al., 2021) in Methods section 2.6 - Comparison: CheZOD score predictions and pLDDT in 17 organisms. The numbers given in columns 3 and 4 were recorded after excluding all proteins with non-extractable pLDDTs (inconsistent table format made the extraction impossible in some cases) and after reducing the data to proteins present in both the AlphaFold2 data, as well as SETH's predictions (see Methods section 2.6 - Comparison: CheZOD score predictions and pLDDT in 17 organisms for more details).

| Organism name | UniProt organism id | Number of proteins after merging with SETH's predictions | Number of residues after merging with SETH's predictions |
|---|---|---|---|
| *Arabidopsis thaliana* | 3702 | 16,134 | 6,973,887 |
| *Caenorhabditis elegans* | 6239 | 4,345 | 1,964,126 |
| *Campylobacter jejuni* | 192222 | - | - |

| | | | |
|---|---|---|---|
| *Candida albicans* | 237561 | 1,027 | 503,966 |
| *Drosophila melanogaster* | 7227 | 3,593 | 1,808,013 |
| *Enterococcus faecium* | 1352 | 29 | 11,721 |
| *Escherichia coli* | 83333 | 4,484 | 1,361,025 |
| *Haemophilus influenzae* | 71421 | 1,661 | 505,987 |
| *Helicobacter pylori* | 85962 | 605 | 193,794 |
| *Homo sapiens* | 9606 | 20,108 | 9,216,997 |
| *Methanocaldococcus jannaschii* | 243232 | 1,773 | 490,308 |
| *Mus musculus* | 10090 | 16,839 | 7,988,435 |
| *Mycobacterium leprae* | 272631 | 669 | 236,327 |
| *Plasmodium falciparum* | 36329 | 5,181 | 2,654,087 |
| *Saccharomyces cerevisiae* | 559292 | 6,693 | 2,759,115 |
| *Shigella dysenteriae* | 300267 | - | - |
| *Staphylococcus aureus* | 93061 | 799 | 256,490 |
| *Leishmania infantum* | 5671 | 7,900 | 4,024,174 |
| *Schistosoma mansoni* | 6183 | 13,826 | 6,433,264 |

**Table S3: Parameter numbers for our models.** Shown are the parameter counts of all models trained in our work: (1) LinReg, a linear regression model trained on 1024-dimensional ProtT5 (Elnaggar et al., 2021) embeddings, (2) LinReg1D, a simplification of LinReg, trained only on the most important embedding dimension for disorder prediction, dimension 295, (3) LogReg, a logistic regression model trained on 1024-dimensional ProtT5 embeddings, (4) ANN, an artificial neural network trained on 1024-dimensional ProtT5 embeddings and (5) SETH, a convolutional neural network trained on 1024-dimensional ProtT5 embeddings. All models apart from LogReg were trained to predict continuous disorder as defined by chemical shift Z-scores (CheZOD scores). LogReg was trained to predict binary disorder/order (CheZOD scores binarized with a threshold of 8, disorder: CheZOD score≤8, order: CheZOD score>8; (Nielsen and Mulder, 2016)).

| Model | Number of parameters |
|---|---|
| LinReg | 1025 (1 intercept) |
| LinReg1D | 2 (1 intercept) |
| LogReg | 1025 (1 intercept) |
| ANN | 3087 |
| SETH | 143,529 |

**Table S4: Full organism names for abbreviations of Fig. 6.** Organism names from Swiss-Prot (The UniProt et al., 2021), which were shortened when showing that SETH’s predictions capture evolution (Fig. 6). The abbreviations with their full names are shown in alphabetical order in each group.

| Group | Abbreviation | Full name |
|---|---|---|
| Eucaryota: Metazoa | Cattle | *Bos taurus* |
| | C. elegans | *Caenorhabditis elegans* |
| | D. rerio | *Danio rerio* |
| | Drosophila | *Drosophila melanogaster* |
| | G. gallus | *Gallus gallus* |
| | Human | *Homo sapiens* |
| | Mouse | *Mus musculus* |

| | | |
|---|---|---|
| | Orangutan | *Pongo abelii* |
| | Rat | *Rattus norvegicus* |
| | X. laevis | *Xenopus laevis* |
| | X. tropicalis | *Xenopus tropicalis* |
| Eucaryota: Fungi | S. cerevisae | *Saccharomyces cerevisiae* |
| | S. pombe | *Schizosaccharomyces pombe* |
| Eucaryota: Viridiplantae | Arabidopsis | *Arabidopsis thaliana* |
| | Rice | *Oryza sativa subsp. Japonica* |
| Bacteria: Actinobacteria | M. bovis | *Mycobacterium bovis* |
| | M. tuberculosis | *Mycobacterium tuberculosis* |
| Bacteria: Proteobacteria | B. mallei | *Burkholderia mallei* |
| | E. coli | *Escherichia coli* |
| | H. influenzae | *Haemophilus influenzae* |
| | H. pylori | *Helicobacter pylori* |
| | P. aeruginosa | *Pseudomonas aeruginosa* |
| | R. palustris | *Rhodopseudomonas palustris* |
| | S. paratyphi A | *Salmonella paratyphi A* |
| | S. typhimurium | *Salmonella typhimurium* |
| | V. cholerae | *Vibrio cholerae serotype O1* |
| | Y. pestis | *Yersinia pestis* |

| | | |
|---|---|---|
| Bacteria: Cyanobacteria | P. marinus | *Prochlorococcus marinus* |
| Bacteria: Firmicutes | B. anthracis | *Bacillus anthracis* |
| | B. cereus | *Bacillus cereus* |
| | B. subtilis | *Bacillus subtilis* |
| | C. botulinum | *Clostridium botulinum* |
| | S. aureus | *Staphylococcus aureus* |
| | S. epidermidis | *Staphylococcus epidermidis* |
| | S. pneumoniae | *Streptococcus pneumoniae* |
| Archaea | M. jannaschii | *Methanocaldococcus jannaschii* |

**Table S5: Residual similarities between train and test set.** The table shows the results of running MMSeqs2 (Steinegger and Söding, 2017) easy-search on the CheZOD117 set against the redundancy reduced training set CheZOD1174 (see Methods 2.1) with --min-seq-id 0.2, -s 7.5, --cov-mode 0 (=bi-directional coverage) and various coverage (-c) thresholds. The sequence identity between 18446 and the training protein 16173 was 0.33 at an alignment length of 61. The sequence identity between 6580 and the training protein 6503 was 0.31 at an alignment length of 87 and lastly, the sequence identity between 11388 and the training protein 4334, was 0.26 at an alignment length of 82.

| Coverage threshold | Sequences of the test set with alignments to training proteins |
|---|---|
| 0.7 | - |
| 0.6 | - |
| 0.5 | 6580, 11388 |
| 0.4 | 6580, 11388 |
| 0.3 | 18446, 6580, 11388 |

| | |
|---|---|
| 0.2 | 18446, 6580, 11388 |
| 0.1 | 18446, 6580, 11388 |
